# Homophilic wiring principles underpin neuronal network topology *in vitro*

**DOI:** 10.1101/2022.03.09.483605

**Authors:** Danyal Akarca, Alexander W. E. Dunn, Philipp J. Hornauer, Silvia Ronchi, Michele Fiscella, Congwei Wang, Marco Terrigno, Ravi Jagasia, Petra E. Vértes, Susanna B. Mierau, Ole Paulsen, Stephen J. Eglen, Andreas Hierlemann, Duncan E. Astle, Manuel Schröter

## Abstract

Economic efficiency has been a popular explanation for how networks self-organize within the developing nervous system. However, the precise nature of the economic negotiations governing this putative organizational principle remains unclear. Here, we address this question further by combining large-scale electrophysiological recordings, to characterize the functional connectivity of developing neuronal networks *in vitro*, with a generative modeling approach capable of simulating network formation. We find that the best fitting model uses a homophilic generative wiring principle in which neurons form connections to other neurons which are spatially proximal and have similar connectivity patterns to themselves. Homophilic generative models outperform more canonical models in which neurons wire depending upon their spatial proximity either alone or in combination with the extent of their local connectivity. This homophily-based mechanism for neuronal network emergence accounts for a wide range of observations that are described, but not sufficiently explained, by traditional analyses of network topology. Using rodent and human monolayer and organoid cultures, we show that homophilic generative mechanisms can accurately recapitulate the topology of emerging cellular functional connectivity, representing an important wiring principle and determining factor of neuronal network formation *in vitro*.

## INTRODUCTION

During mammalian brain development, neuronal networks demonstrate remarkable selforganization that gives rise to complex topological properties, including a greater-than-random clustering and modular structure^1,2^, non-random occurrence of specific network motifs^3,4^, hierarchies^5^, heavy-tailed connectivity distributions^6,7^, and richly interconnected hubs^8,9^. Studies have indicated that these distinctive characteristics likely endow neuronal networks with robustness and the capability to support dynamic functional computations^10,11^, however, our understanding regarding the underlying mechanisms and wiring rules that give rise to these features is still incomplete.

Neuronal network development can be characterized across spatial scales^12,13^. At the cellular level, neurons form computational units within circuits. Here, the role of individual neurons can be determined by a combination of factors, such as their laminar location, connectivity, neurochemical sensitivities and morphology^14^. During embryonic development, a series of spatiotemporally defined genetic and activity-dependent programs^15^ regulate the expression of cell-type specific recognition molecules to initiate axonal and dendritic outgrowth, which ultimately leads to the formation of synapses^16–18^. Although there is now a large body of evidence on the mechanisms of specific guidance cues during circuit formation^19^, linking this knowledge to explain the emergence of complex topological features remains challenging.

At the whole-brain level in humans^20^, connectivity between brain regions can be inferred via diffusion tensor imaging (DTI)^21,22^, as myelinated axonal connections, or functional magnetic resonance imaging (fMRI)^23^, as correlated patterns of activity. Inferring connectomes from fetal brains *in utero^24^*, or from preterm infants^25^, have confirmed the early presence of organizational hallmarks, such as hubs, a rich-club architecture and a modular small-world organization. Building on such architectures, studies demonstrate that the functional role and organization of brain regions later on is shaped by their inter-regional connectivity and that a region’s inputs during development influences its functional specialization^26^. This principle allows brain regions to undergo a spatially-organized functional shift, for example, from distinct sensory and motor systems to more integrated connections with association cortices, likely supporting an acceleration in cognitive development^27^.

There is growing evidence that some key organizational properties of nervous systems are conserved across scales, and, in some cases, across species^12,13,28–32^. Nervous systems both at the macro- and micro-scale, for example, have been shown to entail a canonical pattern of small-worldness^33,34^, a rich-club topology^35–37^ and a modular structure^1,38^. These complex organizational hallmarks allow for functional hierarchies, in which distinct segregated modules perform specialized local computations. While the later may reflect basic representational features of incoming signals, intermediary nodes integrate those signals to code for a more complex representation of the incoming signals^39^.

One prominent explanation for these consistent organizational hallmarks is that they reflect the economics of forming and maintaining connections^40–42^. Given finite available resources, trade-offs between incurred costs (e.g., material, metabolic) and functional value have to be made continually by all distributed units to ensure optimal network function. In this view, ideas, such as Peters’ rule^43^, which suggest that synaptic contacts simply occur if neurons are close enough in space and if their axons and dendrites overlap, are not sufficient to explain the existence of a specific connection^44–47^. Although spatial embedding and neuron morphology clearly have an impact on cortical network architecture^48,49^, costly features, such as long-range connectivity hub cells or regions, likely exist because they confer some additive functional value that outweighs the cost of its formation and maintenance^40^. Such principles likely apply not only to the connectivity between brain regions, but also to the cellular and subcellular level^10,20,49,50^.

If an economic trade-off represents an important principle that guides network development, then it is important to consider the specific mechanisms that determine the outcome of this trade-off. Advances in generative network models (GNMs) provide a formal way of testing competing mechanistic accounts of network formation^51–63^. These computational models simulate the probabilistic formation of networks over time under specific mathematical rules. For example, recent whole-brain DTI work has shown that structural inter-regional connectivity can be simulated with a GNM^51^ which uses a simple economic wiring equation balancing connection costs with topological value^51,53,61^. These two components together define the probability of connections forming iteratively over time. However, the extent to which such models reflect underlying biological processes remains unclear due to the indirect nature of *in vivo* imaging.

In the present study, we test whether key economic trade-off rules are also conserved at the cellular scale. Analyses are carried out on spike-sorted, high-density microelectrode array (HD-MEA) recordings that allow us to directly record from individual neurons and track both their activity and connectivity across development^64–66^. Previous studies have quantified the functional couplings among neurons during spontaneous electrical activity and suggested that the local topological statistics are related to firing properties that may drive neuronal self-organization^67–69^. Here, we expand on these analyses and use functional connectivity inferred for individual neurons tracked over several weeks to probe how spiking patterns of neurons facilitate the implementation of economic wiring. Moreover, we translate prior GNM research at the level of inter-regional brain connectivity^51–53,56,58–61,63^ to the cellular scale, to test whether common generative wiring principles are recapitulated *in vitro*.

We acquire and analyze HD-MEA network recordings from populations of developing primary cells (PCs) derived from dissociated embryonic rodent cortices, three different lines of purified human induced pluripotent stem cell (iPSC)-derived neurons and sliced human embryonic stem cell (hESC)-derived cerebral organoids (hCOs). We compare the performance of different GNMs, and probe whether they can account for the emerging network topology. We also examine the effect of neuronal plating density on network topology and test whether GNMs are capable of recapitulating the local organizational properties of the observed networks. Moreover, by chronically blocking GABA_A_ receptors, we probe how GABAergic signaling impacts neuronal variability and the subsequent formation of connections in the network. Across four model types, comprising 13 wiring rules, we find that homophilic wiring reliably recapitulates the topology and developmental trajectory of neuronal networks at the cellular scale. Homophily may therefore represent an important wiring principle in which local network structure is refined via activity-dependent mechanisms.

## RESULTS

### Tracking developing neuronal networks at single-cell resolution

The datasets of this study comprise recordings of rodent and human neuronal networks that were plated and maintained on high-density multielectrode arrays (HD-MEAs)^64^ as previously described (**Figure 1a**)^65,70,71^. Our primary analysis focuses on primary rat embryonic day 18/19 cortical cultures (PCs), which were plated at two different plating densities (sparse cultures: 50,000 cells per well, n=6 cultures; dense cultures: 100,000 cells per well, n=12 cultures) and used to follow neuronal network development across several weeks *in vitro*. **Figure 1** provides an overview on the experimental pipeline. We also analyzed human induced pluripotent stem cell (iPSC)-derived neuron/astrocyte co-cultures, containing predominantly glutamatergic, dopaminergic and motor neurons (plated at a density of 100,000 cells per array), as well as, sliced human cerebral organoids (hCOs; n=6 slices). For the complete details of the datasets analyzed in the study, see **Methods; *Rodent primary cortical neuronal cultures; Human induced pluripotent stem cell-derived neuronal cultures; Human cerebral organoid slice cultures***. All summary statistics across datasets are provided in **Supplementary Table 1**.

**Figure 1.**
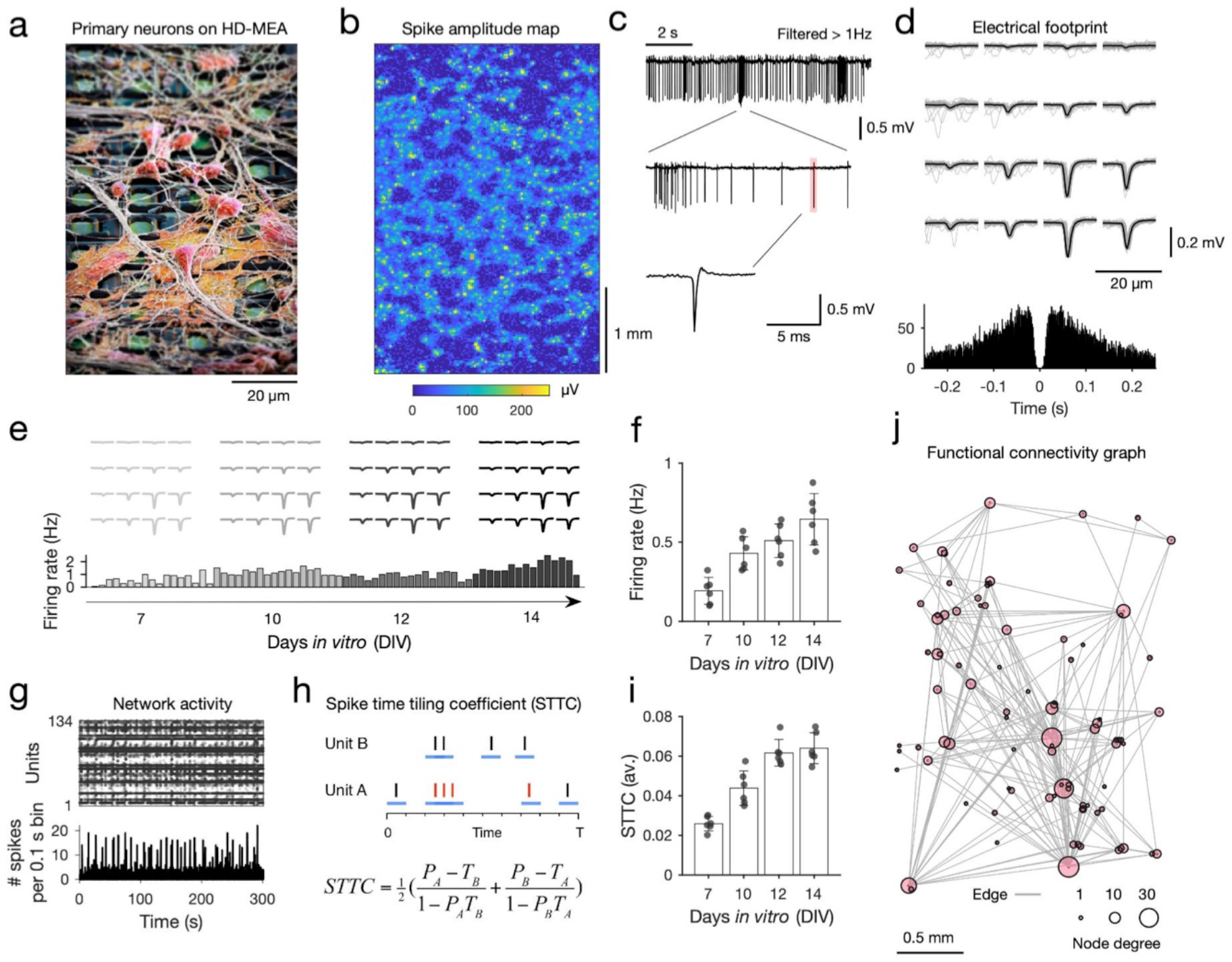
High-density microelectrode array extracellular recordings of developing neuronal cultures. **a** Scanning electron microscope (SEM) image of primary neurons plated on a high-density microelectrode array (HD-MEA; neurons are colored in red, electrodes in green; the electrode center-to-center distance is 17.5 μm). **b** Activity scan across the entire HD-MEA for an example sparse PC neuron culture at DIV 14. Colors depict the average amplitude of online-detected multi-unit activity per electrode (yellow colors indicate the location of potential neurons, i.e., high amplitude deflection, on the array; the array dimensions are 120 x 220 electrodes; electrode amplitude values are averaged over 1 min recordings). **c** Example extracellular trace recorded from one electrode (10 s), with high action potential (AP) spiking activity and bursts (middle panel); a single AP is depicted in the lower panel. **d** Electrical footprint (EF) of a single spike-sorted unit on the HD-MEA. The EF is the AP-triggered extracellular potential signature of a single unit on the HD-MEA, here depicted across 16 electrodes of a high-density recording block (in light gray: 100 AP-triggered cutouts for this EF; in black: the average EF. The lower panel shows a spike autocorrelogram for this unit. **e** Tracking of individual EFs across development *in vitro*. The upper panel depicts the EFs inferred for the four recording time points (DIV7, 10, 12, 14); the lower panel shows the average activity of the tracked unit (bin size: 100 s; gray to black colors correspond to the four recording time points; 30 min per recording day). **f** Spontaneous electrical activity of neuronal cultures, their average firing rate, increased significantly with development (n=6; sparse PC networks). **g** A spike raster plot shows the spike-sorted activity for one culture (300 s-long zoom in on a network recording with 134 units); the lower panel depicts the binned activity of the same recording (bin size: 0.1 s). All neuronal networks showed a mixture of regular and more synchronized activity periods (network bursts). **h** In order to probe neuronal network development, we used the spike time tiling coefficient (STTC) to infer functional connectivity statistically. The four parameters required to calculate STTC connectivity graphs between Unit A and Unit B are P_A_, P_B_, T_A_, and T_B_. T_A_ is calculated as the fraction of the total recording time ‘tiled’ by ±Δ*t* of any spike from Unit A. P_A_ is the proportion of spikes from Unit A, which lies within ±Δ*t* of any spike(s) of Unit B (spikes in red). The spike trains for Unit A and B are depicted as black bars; ±Δ*t* is depicted as blue lines for each spike. The significance of each STTC value was estimated against jittered surrogate spike train data. **i** The average STTC increased significantly with development (n=6; sparse PC networks). **j** Example functional connectivity graph inferred from a DIV 14 PC network (only the top 2% strongest connections are displayed; each dot represents the physical location of a putative neuron on the HD-MEA; the dot size corresponds to the nodal degree of the neuron.

To record the emerging spontaneous neuronal activity of neuronal cultures and to track developing rodent PC neuronal networks at single-cell resolution, we first acquired whole-array activity scans to localize the neurons (**Figure 1b**), and then selected up to 1024 readout electrodes, configured into 4×4 electrode high-density blocks (**Figure 1d, e**), at the respective recording start points (i.e., days *in vitro* (DIV) 7 for the sparse PCs and DIV14 for the dense PCs networks). Recordings with the same electrode configuration were acquired at the consecutive recording time points and concatenated for spike sorting^72^. Spike-sorted network data enabled us to assign the extracellular electrical activity to individual neurons and to follow them across development (**Figure 1e**; see also **Methods; *Spike-sorting and post-processing***; number of neurons tracked for sparse PC networks: 130±28 (mean±S.D); number of neurons tracked for dense PC networks: 115±27; **Supplemental Figure 1**). In line with previous works, we find that PC neuronal networks developed robust network burst activity (**Figure 1c, g**) and that the firing rate of tracked units increased significantly in the first weeks of development (repeated measures analysis of variance (rmANOVA): F(3,12)=7.02, p=5.62×10^-3^; n=6 sparse PC networks; **Figure 1f**).

To infer functional connectivity between neurons statistically and to characterize neuronal network development of tracked neurons over time, we computed the Spike Time Tiling Coefficient (STTC) among all neurons above a minimum firing rate threshold (0.01 Hz; **Figure 1h-j**)^73^ and compared inferred empirical values to jittered surrogate values (see **Methods; Functional connectivity inference**). We use the STTC because it controls for the variability in firing rates between neuronal units and culture types. STTC values primarily reflect short-latency co-activity rather than firing rates *per se*^73^. Although considered a robust measure of pairwise correlations between spike trains, it is important to note that the connectivity graphs inferred by the STTC do not necessarily match the underlying synaptic connections^74^. We also apply a recently published transfer entropy algorithm to show that the main results of this study translate to other measures of (effective) connectivity^75^. In line with previous research^76,77^, we find that overall STTC values increased with development (F(3,12)=11.82, p=6.77×10^-4^, sparse cultures; **Figure 1i**), as did the network density of inferred functional connectivity graphs (F(3,12)=11.08, p=8.97×10^-4^, n=6 sparse PC networks; **Supplementary Figure 2a**). The probability of inferred STTC connections decayed with inter-neuronal distance (**Supplemental Figure 2b**).

### Generative network models of functional neuronal networks in vitro

Following STTC connectivity inference, we set out to describe the topology of these networks using graph theory, which provides a mathematical framework for capturing the topological properties of each node within the network, and the network as a whole. In **Figure 2a** we highlight three common topological measures (nodal degree, clustering, and betweenness centrality) and one geometric measure (edge length) that we will use throughout the current paper (see **Methods; *Network statistics*** for more details^9,78^). In **Figure 2b** we show how these statistics allowed us to compute the node-wise statistics for each functional connectivity graph, and to establish the distribution of different statistics across the network.

**Figure 2.**
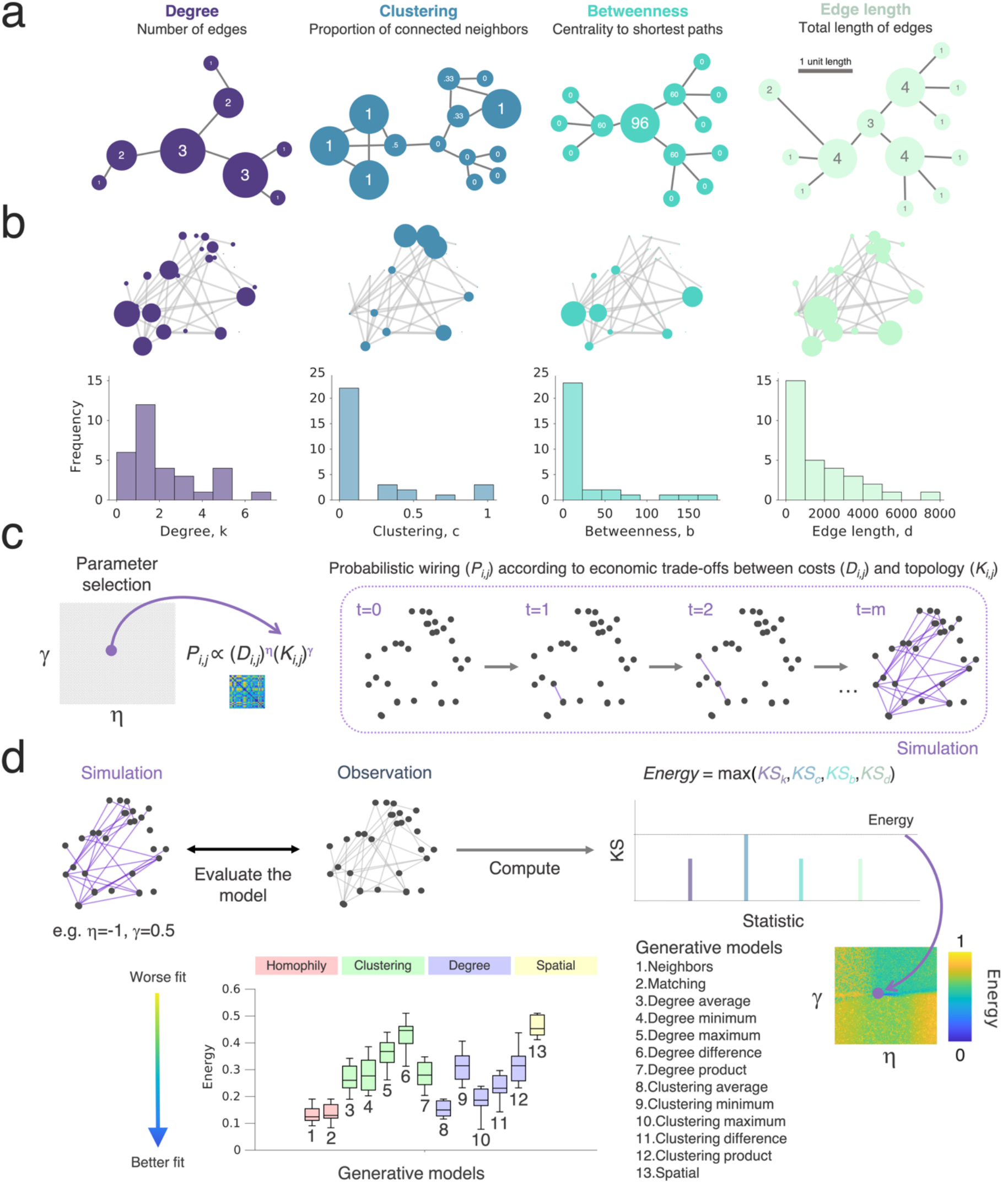
Probing wiring principles via generative network models. **a** Example networks highlighting four common graph theoretical metrics. For each schematic, the node size corresponds to the respective graph statistic, and we provide the statistic within the node. The panel on the left shows a schematic for the degree (blue), which relates to the total number of edges that each node has. The left-middle panel shows the clustering coefficient (green), which is a measure of how nodes in a graph tend to cluster together; the right-middle panel shows the betweenness centrality (pink), which relates to the number of shortest paths in the network which pass through the node; the panel on the right shows a schematic for the total edge length (yellow), which relates to the total sum of the edge lengths among all its connections for each node. **b** Functional connectivity graphs, as inferred from HD-MEA network recordings, were characterized by graph theoretical means. Each panel on the top row shows the same network, with the node size corresponding to the respective graph statistic (degree, clustering, betweenness centrality, total edge length). The lower row shows histograms with the distribution of the metric across the network. **c** The generative network model works by simulating network development probabilistically according to the costs incurred in the network (here reflected by the *D_i,j_* term, which is modeled as the Euclidean distances) and the topological values. Both terms are parameterized, meaning that for each simulation there is an altered tuning in how costs and values influence the probability with which nodes form connections iteratively. Parameters (η and γ) alter direction and magnitude to which the costs and values respectively influence wiring probabilities. For any combination of wiring parameters, we simulate a network developing over time until it reaches the number of connections, m, equal to the observed empirical network. **d** Comparing the simulated and empirically observed network, we fit the generative network model to achieve a simulation that is the least dissimilar to the observation. This is quantified by the energy equation, and is shown by dark blue in the parameter space. The energy equation is given by the maximum KS statistic across degree, clustering, betweenness centrality and edge length. Each parameter combination is plotted on an energy landscape, which demarcates the model fits across the two-dimensional parameter space. Lower energy values correspond to better fits.

Although graph theoretic measures provide a way to mathematically formalize the topology of networks, they do not provide an explanation as to what topological attachment principles may have shaped network development. To do this, we tested 13 wiring rules that may best explain the self-organization of cellular-level functional connectivity graphs over time. Each of these 13 rules that we tested is given in **Supplementary Table 2.** To simplify, for the main report we group the 13 rules into four broader attachment mechanisms by which they work: (i) *Homophily*, where a neuron, *i*, preferentially wires with another neuron, *j*, as a function of the similarity between *i* and *j* in terms of the other neurons they connect to (we expand on this later). (ii) *Clustering*, where neuron *i* preferentially wires with neuron *j* as a function of neuron *i* and *j’s* independently computed clustered connectivity or (iii) *Degree*, the number of their connections. (iv) *Spatial*, where neurons only wire with other neurons as a function of the physical distance between each other.

We start with a model for which *spatial proximity* is the exclusive determining factor for the emergence of a connection. If neurons connect according to Ramón y Cajal’s laws of conservation^79^, balancing the costs of connections with the functional benefits that they provide, one may predict neurons would simply connect to their geometrically closest neighbors to minimize the cost of maintaining connections^13,40,79^. However, other network models incorporating the principle of preferential attachment, such as the Barabási-Albert model^80^, suggest that a rich-get-richer principle may drive the emergence of topology. This is where the more connections a neuron has, the greater the probability of forming more connections thus leading to scale-free, power law degree distributions^62,80^. Therefore, we also apply *degree-based models*, where connections are more likely the more connections one or both neurons have. Another canonical network model, the Watts–Strogatz model^81^, illustrates how networks deal with the tradeoff between local and global processing—nodes form clusters at the cost of global integration, though local topology is key for modular structure and regional functional specialization, hence are a key component of small-world networks^81^. We therefore include *clustering-based models*. However, clustering-based rules compute how many neighbors of neuron *i* are connected to each other and how many of neuron *j*’s neighbors are connected to each other, separately—they are not necessarily part of the same cluster with the same neighbors. Thus, the benefit of a connection between *i* and *j* may be less than if they had similar neighbors. This could provide a mechanism for neurons with similar functional purposes to connect. Hence, we also include *homophily-based models* based on similarity in neighborhoods between neuron pairs.

We undertook our simulations using a generative network model, which was previously used to probe whole-brain network organization^51,53,61^. Generative network models develop *in silico* networks according to an economic trade-off, in which new connections are iteratively formed depending on both the modeled costs and values (**Figure 2c**). The generative algorithm is expressed as a simple wiring equation^56,61^, which is updated over time:

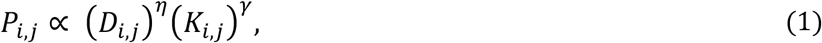

where the *D_i,j_* term represents the “costs” incurred between neurons modeled as the Euclidean distance between tracked neurons *i* and *j* (**Supplementary Figure 3**). The *K_i,j_*, term represents how single neurons *i* and *j* “value” each other, given by an arbitrary topological relationship which is postulated *a priori* (also termed, “wiring rule” given mathematically in **Supplementary Table 2**). *P_i,j_* reflects the probability of forming a fixed binary connection between neurons *i* and *j*. This is proportional to the parametrized multiplication of costs and values. Two wiring parameters, η and γ, respectively parameterize the costs and value terms, which calibrate the relative influence of the two terms over time. We detail the generative network algorithms used in **Methods; *Generative network model***. By iterating through different *K_i,j_* terms and wiring parameter combinations, we can formally assess how variations in the generative model give rise to synthetic networks, which are statistically similar to those experimentally observed (**Figure 2d**). To assess this similarity, the first test comes in the form of an energy equation^53^, which computes the Kolmogorov-Smirnov (KS) distance between the observed and simulated distributions of individual network statistics. It then takes the maximum of the four KS statistics considered so that, for any one simulation, no KS statistic is greater than the energy:

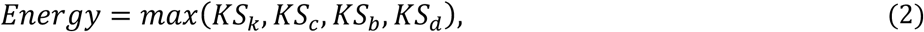

where *KS* is the Kolmogorov-Smirnov statistic between the observed and simulated networks at the particular η and γ combination used to construct the simulation, defined by the network degree k, clustering coefficient c, betweenness centrality b and Euclidean edge length d. These four measures are critical statistical properties of realistic networks, have been used in prior GNM research to benchmark model fits^51,53,54,61,63^, and have featured within well-documented simulated network models^80–82^.

For each empirical network, we simulated 20,000 networks across a wide parameter space (with η and γ limits −10 to 10) across the 13 wiring rules, for each network across all available time points. In the present study, we used this wide parameter space as there is little prior work guiding our choice of parameters; we also did not select a seed network. A Voronoi tessellation procedure^53^ was used as the parameter selection algorithm (see Methods; Parameter selection for details).

### Homophilic wiring principles recapitulate the topology of developing rodent neuronal networks in vitro

Previous studies employing generative models of human macroscopic structural brain organization have shown that generative rules based on homophilic attachment mechanisms can achieve very good model fits^51,53,60,61,63^. Homophily-based wiring prioritizes the wiring of nodes preferentially to those with overlapping connectivity patterns (e.g., via neighborhoods or connectivity profiles). For example, under a matching generative model^51,53^, if two nodes have a large proportion of their connections with the same nodes, they will have a correspondingly high matching score because they have similar connectivity profiles. This matching score is *homophilic*, because the measure is defined in terms of similarity (the Greek *homós*, “same”) and preference (*philia*, “liking”). To test what generative models can best simulate microscale connectivity, we applied the generative procedure to inferred STTC functional connectivity graphs.

We first investigate the sparse (50,000 cells per well) PC rodent networks at DIV 7, 10, 12 and 14. As previously shown (**Figure 1**), PC rodent networks underwent significant developmental changes during this time period, and this is reflected by large topological changes in their functional networks (**Supplementary Figure 2**). We find that over the developmental time course, generative models utilizing the homophilic attraction principle as their generative mechanism produce networks with increasingly good model fits (the energy) compared to the degree-, clustering- and spatially based rules tested (**Figure 3a**). By DIV 14, homophily alone performs best (p<4.11×10^-5^ for all pairwise comparisons and Cohen’s *d*>1.46 reflecting a very large effect size; **Supplementary Table 3** shows all statistical comparisons). The single best performing homophily model, according to the energy equation, was the ‘matching model’ (see **Supplementary Table 2** for detail), which generates network topology according to the overlapping connectivity between nodes (**Supplementary Figure 4**). To assess the extent to which these findings go beyond what may be expected by chance, we further examined model performances across a range of alternative procedures for formalizing the generative models, including a *K_i,j_* only formalization (where space has no influence), comparison to density- and size-matched random null models, and a comparison of our networks to those constructed using transfer entropy (a model-free effective connectivity measure) rather than the STTC. We find that, regardless of the model specification or connectivity measure tested here, homophily performs best when approximating empirical networks, but not randomized networks (**Supplementary Figure 5**). In **Figure 3b**, we also show an immunohistochemistry staining of an example PC rodent network at DIV 14 and **Figure 3c** shows the energy landscapes acquired from the matching generative model at this same timepoint.

**Figure 3.**
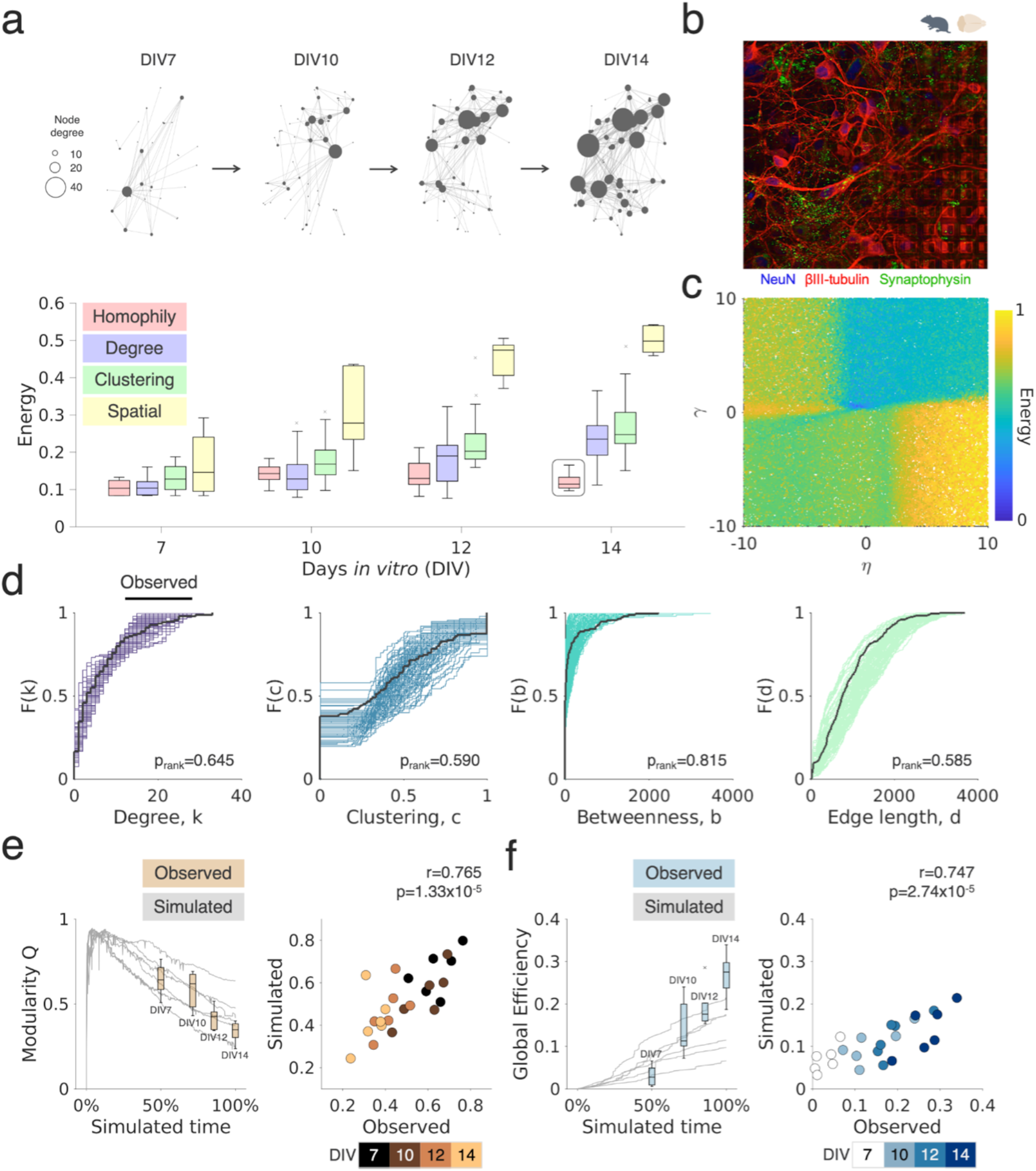
Homophilic principles underpin rodent primary network development and explain variance in their maturational trajectories. **a** The top row shows a representative rodent primary cortical (PC) network developing over the first two weeks *in vitro* on the HD-MEA (DIVs 7, 10, 12 and 14; sparse neuron plating density; gray nodes correspond to single neurons; the size of each neuron corresponds to its nodal degree). Below we show the energy of the top performing simulated networks, given across the four main categories of studied wiring rules. Boxplots represent the median and interquartile range (IQR); outliers are demarcated as small black crosses, and are those which exceed 1.5x the IQR away from the top or bottom of the box. **b** Immunohistochemical staining of a rodent PC culture (control experiment; NeuN staining in blue, βIII-tubulin in red, and Synaptophysin in green). **c** All energy landscapes of the matching generative network model for DIV 14. **d** Cumulative density functions (CDFs) of the matching generative model, showing that simulated and observed statistics overlap very well. CDFs are shown across the four network statistics included in the energy equation of the top=99 simulations for the matching model (best performing generative model) compared to an observed network (black line). Subsequent p_rank_ values were computed using a Monte Carlo sampling procedure. **e** and **f** Simulated network trajectories were examined to determine, if the later generative model’s trajectory (DIV 14) recapitulated earlier observed statistics. Simulated network statistics were derived by computing the modularity Q statistics (**e**) and global efficiency (**f**), at each time-point, within the generative model that was best fit to the DIV 14 networks. Results were scaled to the developmental time so that observations and simulations could be compared directly.

The matching generative model, beyond providing the lowest energy values (i.e., very good model fits compared to other models), also produced synthetic networks whose aggregate nodal distributions were statistically indistinguishable from the experimentally observed networks (**Figure 3d**). We formally demonstrated this using a Monte Carlo sampling procedure^83^ to directly compare the statistics produced by the well-performing simulations with the observations (degree, p_rank_ =0.645; clustering, p_rank_ =0.590; betweenness, p_rank_ =0.815; edge length, p_rank_ =0.585; for detail of this procedure, see ***Methods; Cost functions***). In **Supplementary Figure 6** we provide the same bootstrapping analysis, but for each of the best performing generative models of each model class (spatial, clustering average and degree average models). It is of note, that this matching generative model was the only model, that was able to produce statistically indistinguishable results when compared to the experimental observations; the next best performing non-homophily model, the degree average model, failed to approximate both the empirical edge lengths (p_rank_=0.025) and participation coefficients (p_rank_=0.04).

Next, we asked how well generative models would approximate the time-course trajectories of neuronal network formation. An advantage of the GNM approach is that it allows one to decompose the developmental trajectory. Indeed, if networks are developing according to a homophilic attachment principle then the statistical properties of those simulated trajectories should vary in accordance to our longitudinal observations. To test this, we computed and compared the trajectories of two global network measures of segregation (modularity index Q^2^) and integration (global efficiency^84^) across the time-course of sparse rodent PC network development. We used these measures because they were both not included in the energy equation (**Equation 2**)—and it is well established, that they capture fundamental aspects of how efficient information can be processed across networks^85,86^. Next, we selected the best fitting model at DIV 14, and decomposed the simulated trajectories up to that point. This allowed us to test whether these simulated trajectories were consistent with the earlier longitudinal observations at DIV 7, 10 and 12.

To do this, we compared each of the longitudinal observations (DIV 7, 10, 12 and 14) to the simulation at the corresponding time-point of the DIV 14 developing simulation (i.e., DIV 7, 50%; DIV10, 71%; DIV12, 86% and DIV14, 100%). **Figures 3e, f** shows the developmental trajectories of modularity and global efficiency for individual simulations (the gray lines) along with the overlaid observed time-points. Simulations using the homophily generative model clearly captured the same developmental trend for modularity (decreasing over time) and efficiency (increasing over time) and accounted for a substantial amount of variance in both metrics (modularity: R^2^=58.5%, r=0.765, p=1.33×10^-5^; efficiency: R^2^=55.8%, r=0.747, p=2.74×10^-4^).

### Functional topology and generative model outcome hold in both sparsely and densely plated cultures

So far, we quantified STTC connectivity graphs derived from sparse PC neuronal cultures, that is, cultures plated with an initial density of 50,000 cells per well (1000-1500 cells/mm^2^). Of note, the actual numbers of cells on the HD-MEAs are likely smaller due to activity-dependent programmed cell death (apoptosis), that occurs during the first weeks *in vitro*^87,88^. Despite some research on the effect of plating density on the emergence of population activity *in vitro*^89^, synaptic strength and connectivity^90^, there is currently no consensus as to how different plating densities affect neuronal topology. As one critical element of the generative network model is the geometric spacing between neurons, we next probed whether our findings in sparse cultures generalize to networks at higher neuronal plating densities. Therefore, we recorded a second independent dataset of more densely plated rodent PCs (100,000 neurons per well, n=12; see **Methods; *Rodent primary cortical neuronal cultures***) in the exact same way as outlined for the first PC rodent dataset and directly compared both densities at DIV 14.

We found that only the empirical edge lengths and global clustering differed between sparsely versus densely plated PC networks; dense networks showed relatively shorter connections (Mann-Whitney U, p=0.0125) and were more topologically clustered (p=0.0135). All other tested metrics remained very comparable (**Supplementary Figure 7**). The global correlational structure of these statistics also remained stable (**Figure 4a,b**). Given the topological differences in edge lengths and global clustering, we then asked whether this also translated into significant changes in the energy values among the 13 tested generative network models. In **Figure 4c** we show that the model energy is unaffected by plating density (homophily between plating densities, p=0.191). All statistical comparisons, for each time-point in the dense PC networks, are presented in **Supplementary Table 4**; results demonstrate stability not only at DIV 14, but also at later time points (DIV 21 and 28). In **Supplementary Figure 8**, we show the same results, but broken down by each individual model, in addition to showing that this result holds also when considering the average energy over the top 10 and 50 best-performing parameter combinations. In **Figure 4d** we show the energy landscape for both plating densities, which again are very similar.

**Figure 4.**
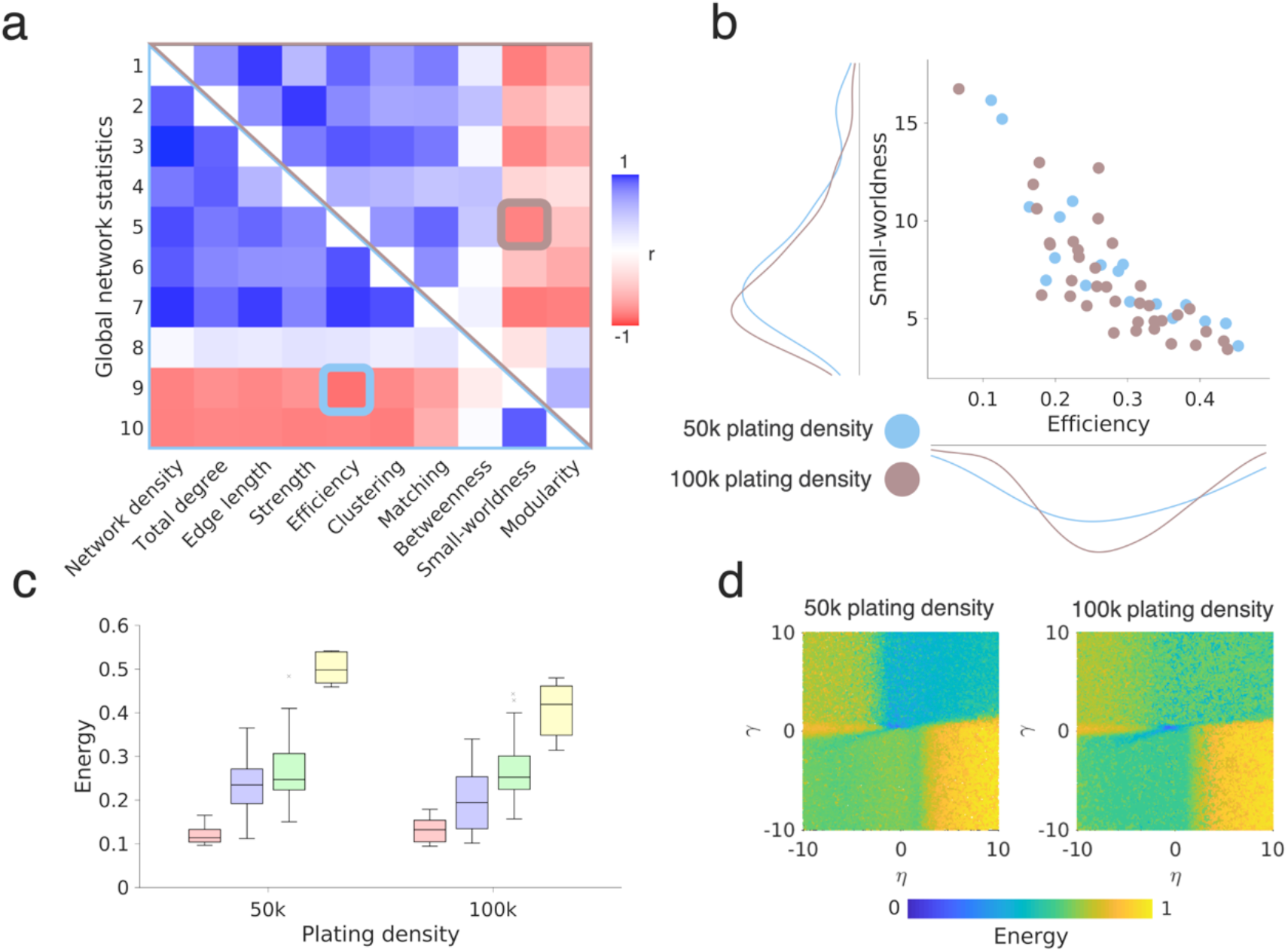
Effect of plating density on network topology and generative network principles. **a** Correlation matrix for global network statistics calculated across sparse (lower triangle) and dense (upper triangle) PC cultures shows a highly similar covariance structure; colors indicate the Pearson’s correlation coefficient. Elements (5,9) and (9,5) of the correlation matrix are further highlighted in panel **b.** This plot shows the correlation between efficiency and small-worldness, for sparse and dense networks (*r*=-0.822, *p*=2.73×10^-14^); each point in the scatter plot corresponds to a single network. **c** Sparse and dense PC networks can be best simulated by the homophily generative models (DIV 14). Note, that the leftmost boxplot is the same as that given in **Figure 3a**. **d** The energy landscapes for the matching generative network model landscapes inferred for the sparse and densely plated PC networks (DIV 14; see also **Figure 3c)**.

### Topological fingerprints arise from homophilic mechanisms in developing neuronal networks in vitro

We have shown that homophily-based generative models produce synthetic networks which are statistically similar to observed functional rodent PCs networks. However, this similarity depends upon the maximum *KS* distance of the four topological statistics as defined in the energy equation. Crucially, this means that while experimentally observed and simulated network statistical distributions mirror each other at the *global* level, the *topological fingerprint* (*TF*) of these network statistics could differ. That is, nodes within simulated and observed networks could have different *local* relationships to one another, because node-wise local organizational properties are not captured *per se* by the existing energy equation. Previous studies have investigated how well generative models can recapitulate local organizational properties and the location of features, such as hub-nodes in the *C. elegans* connectome^91^ or MRI-inferred human brain networks^51,52,60^.

This can be exemplified by the topological relationship between central and peripheral nodes. Nodes which score high in centrality measures (e.g., *betweenness centrality—which* determines how many shortest paths pass through—as shown by the red node in **Figure 5a, left**) tend *not* to sit within segregated modules, meaning it is common that they concurrently score low in measures of segregation (e.g., clustering coefficients, in which neighbors connect to each other). The opposite is true for peripheral nodes (see the green node in **Figure 5a**). This means that when correlating measures of centrality with measures of clustering across a network, the correlation tends to be negligible or negative^39^ (**Figure 5a, right**).

**Figure 5.**
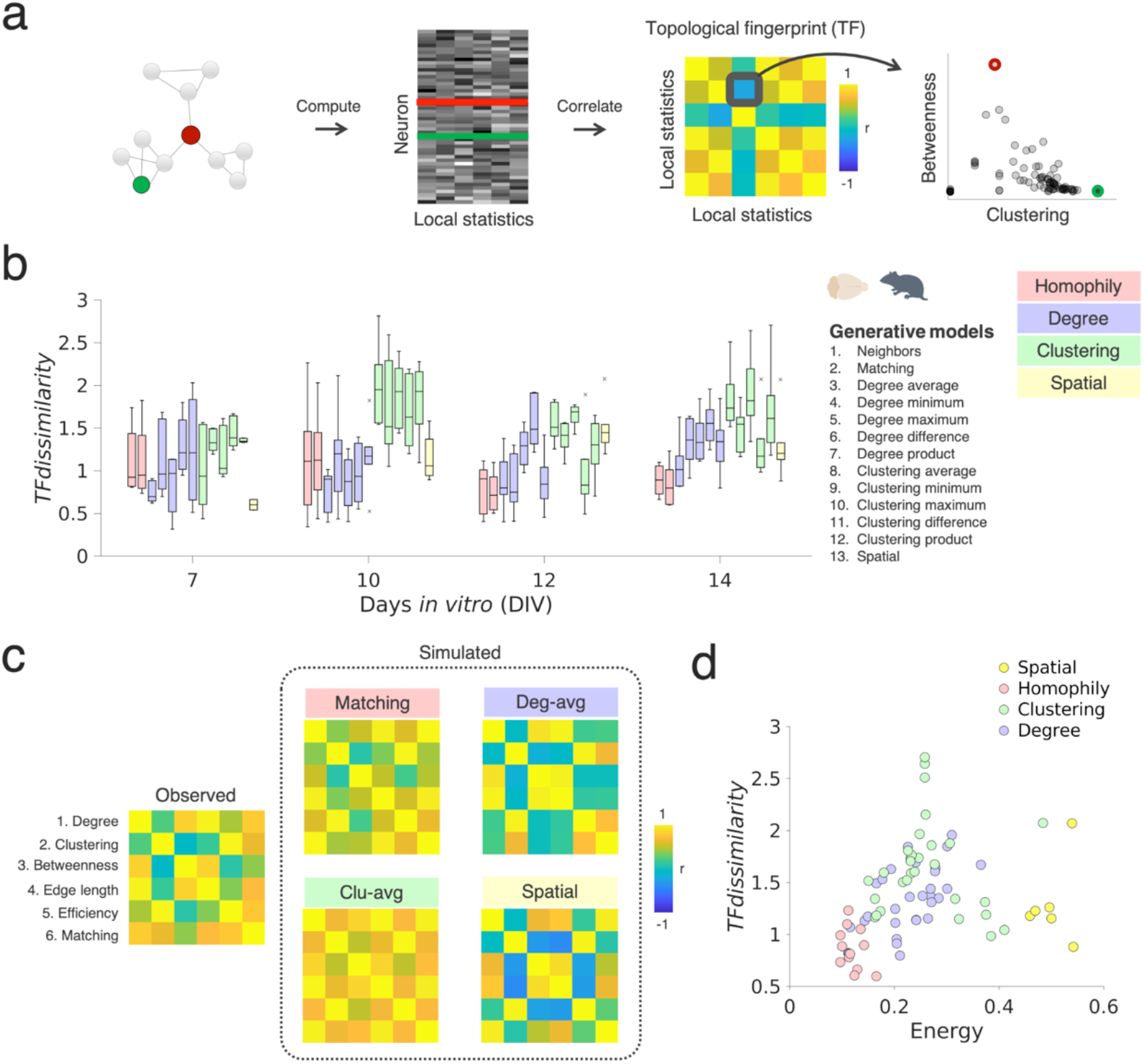
Homophilic generative mechanisms capture the local topology in developing neuronal networks. **a** Schematic illustrating the inference of topological fingerprints (TFs). A TF measure is computed as the Pearson’s correlation matrix among several topological statistics at the nodal level (degree, clustering, betweenness, edge length, efficiency and matching). Each node in **a** corresponds to a single neuronal unit/neuron. The right panel shows the negative correlation between clustering and betweenness, which captures an aspect of the topological structure of the network shown on the left. The color bar is clipped to +/-1 (blue/yellow) for clarity. **b** The *TF_dissimilarity_* measures the extent to which GNM simulations capture the topological structure of the experimentally inferred networks. Homophily generative models, and spatial models, show the lowest *TF_dissimilarity_*, suggesting that both can reconstruct local connectivity patterns in *vitro* (n=6, sparse PC networks; boxplots present the median and IQR; outliers are demarcated as small gray crosses, and are those which exceed 1.5 times the IQR). **c** Averaged *TF* matrix for the empirically observed data (on the left; DIV 14), versus the GNM results obtained from models with the best fits, i.e., the lowest energy values obtained for each model class: the matching rule (top left panel), the clustering-average rule (bottom left panel), the degree-average rule (top right panel) and the spatial mode (bottom right panel). **d** This plot depicts the relationship between energy and *TF_dissimilarity_* values for each sparse PC network broken down by generative rule class (spatial in yellow, homophily in red, clustering in green, degree in blue). Each dot indicates the value of the two model fit functions for a single simulated network.

To assess the ability of generative models to capture these types of local relationships in settings with no anatomical reference space (as neurons are randomly distributed on the HD-MEAs), we provide a very simple cost function, here termed *topological fingerprint dissimilarity* (*TF_dissimilarity_*). The *TF_dissimilarity_* demarcates the ability of *in silico* network simulations to recapitulate observe local hallmarks of organization. It is defined as:

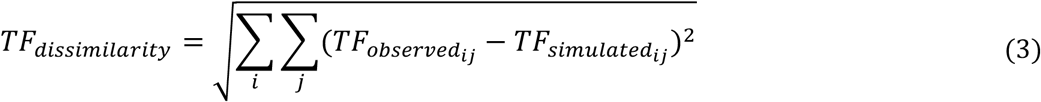

The *TF* is defined by the n-by-n correlation matrix of n local network statistics for the observed network (*TF_observed_*) and its corresponding (simulated) network (*TF_simulated_*). The *TF_dissimilarity_* is subsequently equivalent to the Euclidean norm^92^ of the difference between observed and simulated topological correlation matrices. Here, we use six common measures of topology to compute the *TF* matrix (see **Methods; *Cost functions*** for details). *TF_dissimilarity_* serves as a unitary measure of the difference between the simulated and observed networks in terms of *local* topology, in contrast the energy metric which reflects *global* topology. **Figure 5a** provides a schematic of how *the TFs* were constructed.

If homophily is a plausible attachment mechanism by which single neurons together form networks, we should expect homophily-based GNMs to produce networks with a local topological structure resembling the observed data. To probe this (dis)similarity, we calculated the *TF_dissimilarity_* between each experimentally inferred functional connectivity graph and the best performing simulated network (according to the energy equation), for each of the 13 generative models, and across all recording time points (**Figure 5**).

Results demonstrate that synthetic networks generated with homophilic attachment rules provide the lowest *TF_dissimilarity_* from DIV 12 onwards (**Figure 5b**). These rules also result in the statistically smallest *TF_dissimilarity_* at DIV 14 (compared to degree rules: p=1.91×10^-3^, Cohen’s *d*=1.34; compared to clustering rules: p=1.37×10^-7^, Cohen’s *d*=2.59). Of note, homophily and spatial rules could be distinguished significantly at DIV 12 (p=9.96×10^-4^) but not at other time-points (e.g., at DIV 14; p=0.0952), possibly indicative of spatial costs driving the topological fingerprint at this level of plating density (**Supplementary Table 5**). Interestingly, replicate analyses in the denser PC rodent dataset at DIV 14, 21 and 28 provided almost identical results, but with an even stronger distinguishability of homophilic rules relative to spatial models (**Supplementary Table 6** and **Supplementary Figure 9**), possibly reflecting a weaker preponderance of costs driving topology (mirroring previously found edge length differences; **Supplementary Figure 7**). The left-most panel in **Figure 5c** shows the experimentally observed *TF* matrix averaged over n=6 sparse PC networks; the adjacent panels show average *TF* matrices for the matching, clustering-average, degree-average and spatial generative models. Depicted are the best performing models within their generative rule class.

In sum, our results highlight the importance of assessing GNM simulation performance both in terms of overall global topology (*energy*) and the local topology generated (*TF*). We find that homophily models concurrently outperform the other models on both fronts (**Figure 5d**, see **Supplementary Figure 10** for a replication analysis in the dense rodent PC dataset).

### Effect of GABA_A_ receptor antagonism on network development and dynamics

Previous studies have shown that GABAergic interneurons can act as network hubs and regulate synchronicity of spontaneous activity between neurons that is critical for the formation of connections^93–95^. The role of GABAergic interneurons during development is also relevant for understanding their function in more mature brain circuits^93,94,96,97^. Moreover, alterations in the ratio of GABAergic interneurons and glutamatergic projection neurons, respectively the balance of excitation and inhibition, has been implicated in many neurodevelopmental disorders^98^. Ionotropic GABA_A_ receptors are known to mediate fast inhibitory transmission in the cortex (in contrast to slow inhibition mediated by metabotropic GABA_B_ receptors)^97,99^ and are critical for persistent network activity required for behavioral functions^97^. However, the role played by GABA_A_ receptors in functional network development is not fully understood. Furthermore, absence of GABA could also delay the developmental switch in GABA polarity from depolarizing to hyperpolarizing^100^. Therefore, it is unclear how cellular-scale functional networks would develop in the absence of GABA_A_ receptor-mediated inhibition and whether connections would form on the basis of homophily as seen in our previous results. It is also unknown whether effects of perturbation to GABA_A_ receptor-mediated inhibition transmission on network activity and functional connectivity would be reversible.

To address this, we cultured sparse rodent primary neurons under chronic application of media without gabazine (n=6, control) or with gabazine (n=9 gabazine-treated), a selective GABA_A_ receptor antagonist (see **Methods; *Pharmacological experiments***). Following chronic application of gabazine for two weeks, we washed out gabazine at DIV 14 and performed a final recording at DIV 15 in a subset of the dataset (n=6, washout) to determine the extent to which the cultures recovered. We first examine the differences in spiking dynamics and functional connectivity as a result of GABA_A_ blockade. Next, we examine changes in energy across rules and parameter values within the best-performing rule. We asked whether GABA_A_ receptor blockade affected network formation by a) preventing any connectivity principle being implemented (where all models have high energy), b) altering the connectivity principle being implemented (where a different model than homophily has the lowest energy) or c) altering how homophily is implemented (where homophilic models still have the lowest energy but parameter directions or magnitudes are altered).

The six control and nine gabazine-treated cultures were compared using Mann-Whitney tests; the six gabazine cultures recorded at DIV14 were compared to their respective recordings at DIV15 with Wilcoxon signed rank tests. In line with previous research^88,101^, we find that chronic application of gabazine has a significant impact on single-cell and network firing patterns. Compared to controls, gabazine-treated networks show a more stereotypic burst behavior with less variation in interburst intervals (CV of IBI, Mann-Whitney U: p<0.001; n=6 controls; n=9 gabazine-treated); **Figure 6**). Moreover, we find a trend towards lower firing rates during chronic gabazine treatment at DIV14 (p=0.0567), which was reversed following washout (DIV15; Wilcoxon signed rank: p=0.0156; n=6). Burst rates and the fraction of spikes occurring in bursts were highly variable across gabazine-treated networks and both metrics significantly decreased following washout (p=0.013). We provide a more detailed comparison of the spiking activity between control and gabazine-treated cultures in **Supplementary Figure 11**.

**Figure 6.**
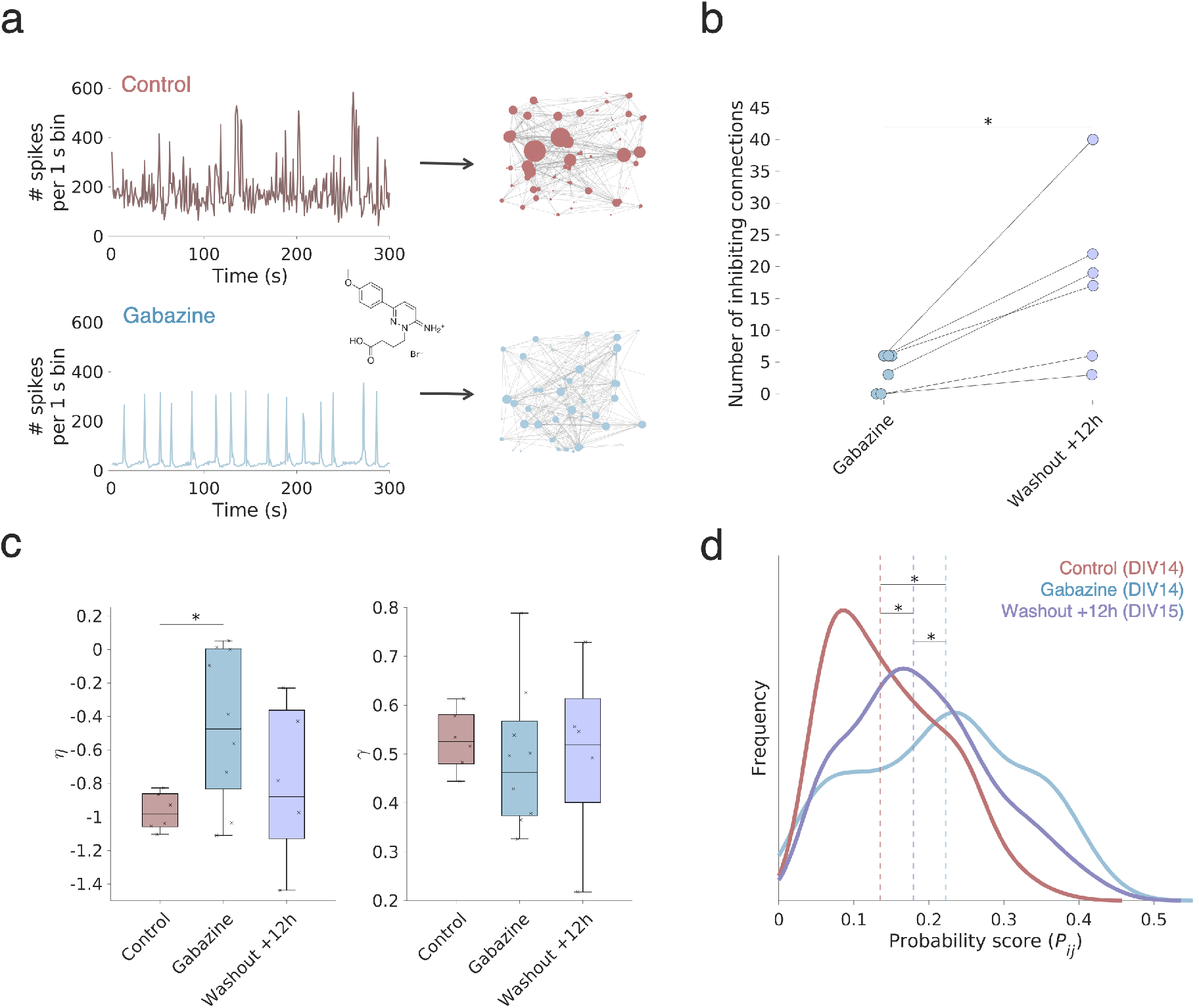
Chronic block of GABA_A_ receptors alters the wiring parameters of the homophily generative model. **a** Representative neuronal population activity of a control/untreated (red, top) and gabazine-treated networks (bottom, blue) at DIV 14. In the gabazine condition, network burst activity is more strongly synchronized, compared to the control networks. **b** the number of putative inhibitory connections increased significantly following the washout of gabazine at DIV 14. Putative pair-wise inhibitory connections were inferred by a cross-correlogram approach and using an algorithm to infer reductions in spike transmission probability^102^. **c** The wiring parameters of the best performing (matching) generative model. Energy values of the top performing simulated networks across control and gabazine-treated cultures are shown in **Supplementary Figure 13a,b**. Of note, the η wiring parameter (left boxplots), which reflects the influence of Euclidean distances on the generative model, decreased in the gabazine condition relative to controls. Wiring parameters were derived from the top n=1 best performing matching simulation. See **Supplementary Figure 13c,d** for a replication across increasing numbers of top performing parameters. **d** The average probability score kernel density distributions (*P_ij_*) for the control (brown), gabazine (cyan) and washout conditions (lilac) recorded in sparse primary culture networks across simulated network development. The average gabazine probability distribution is shifted to the right and flatter compared to the controls and the washout condition. This indicates that wiring becomes more random across all timepoints. In **Supplementary Figure 13e,f** we show all distributions used to construct these averages. Asterisk indicates p < 0.05.

Control and gabazine treated networks differed in global functional connectivity (**Supplementary Figure 12**). On average, gabazine-treated networks showed an increase in average STTC (Mann-Whitney U; p<0.001), a higher network density (p=0.003), and a greater total edge length (p=0.026). Following the washout, average STTC (p=0.031), network density (p=0.031) and edge length (p=0.013) reduced significantly, resembling the untreated cultures. Whereas untreated cultures showed a small fraction of nodes with high nodal strength, indicative of potential hub nodes, we did not see this topological structure during chronic gabazine treatment. That is, control networks had a more positively skewed STTC distribution than controls (p=0.002). This is consistent with the notion that GABA_A_ receptors are involved in regulation of spiking activity and synchrony between neurons in the network, as this was altered by GABA_A_ receptor block.

To further examine the extent to which washing out gabazine returned endogenous inhibitory activity as GABA_A_ receptors are released from blockade, we also computed the spike transmission probability (STP; **Supplementary Figure 13**). This cross-correlogram based metric has been used to infer putative inhibitory functional connectivity from ongoing extracellular spiking activity and to identify neuron pairs with a reduction in spike transmission gain (STG)^102^. Indeed, we find that washout of gabazine led to a significant increase in the number of putative inhibitory connections (Wilcoxon signed rank: p=0.012; **Figure 6b**).

Despite alterations in cellular activity and network topology (**Supplementary Figure 11** and **12**), homophilic generative attachment rules were the best fitting models across both control, gabazine-treated and washout (**Supplementary Figure 14a, b**). However, gabazine cultures exhibit a higher energy relative to controls (p=0.0292; **Figure 6c**), which suggests that homophilic GNMs cannot approximate the topology of gabazine-treated cultures to the same extent as for control cultures. This finding supports our hypothesis that perturbing GABA_A_ receptor-mediated inhibition alters network characterization, even though homophily remains the best-fitting model. After washout of gabazine, however, these same cultures exhibited homophily energy values indistinguishable from controls (p=0.285), which indicates, that this effect may be reversible. **Supplementary Figure 14c** shows how gabazine cultures are best fit when the η homophily wiring parameter increases so that it is closer to zero, i.e., moving from more negative to less negative, decreasing in magnitude (p=0.0360). Lower magnitude η corresponds to a weaker influence of physical distance on the probability of forming connections. On the other hand, γ (which varies the extent to which homophily influences wiring probability) is unaffected (p=0.456). Hence, gabazine weakens the spatial extent of functional connectivity wiring, enabling longer distance connections in the network.

Differences in wiring parameterization may reflect more fundamental differences in neuronal variability elicited via changes in the neural dynamics. Within the homophily generative model, connections form via continual negotiations between the modeled cost and self-similarity that is present between all neurons. If there are clear *relative* winners in this negotiation, for example, connections that are lower cost than all others and connections that are more homophilic than all others, these connections are more likely to form. Of note, the probability of a connection is proportional to the wiring parameter magnitudes (see **Equation 1**). Mathematically, the *P_i,j_* distribution would look like a canonical lognormal distribution that is found at many anatomical and physiological levels of the brain^103,104^, where a small number of possible candidate connections elicit a higher wiring probability while the majority remains low. However, in the example of gabazine, neuronal synchrony increases, which is equivalent to exhibiting less variability in spike times between neurons. Therefore, we hypothesized that this would lead to a flattening of this lognormal distribution, meaning that the resultant topology became more random and spatially distributed. Indeed, in **Figure 6d** we show this to be the case: gabazine-treated networks exhibit a flattened wiring *P_i,j_* distribution relative to both controls (median *P_i,j_* value=0.135 and 0.322 for gabazine & control, respectively; p=1.54×10^-44^, Cohen’s *d*=0.550) and also after gabazine washout (median *P_i,j_* value=0.179; p=5.04×10^-8^, Cohen’s *d*=0.196; see also **Supplementary Figure 14d**). This finding suggests that gabazine alters the network wiring distribution as to become less specific in its wiring, rather than being specific to a smaller number of candidate neurons that are deemed particularly valuable to wire with.

In summary, we find that homophily wiring rules are also the best fitting GNM for neuronal networks chronically treated with gabazine. For the latter, however, homophilic GNMs achieve worse fits compared control cultures. At the level of model parameters, the homophily γ parameter remains stable over all conditions, but it is η - which alters the spatial extent over which wiring is constrained. Interestingly, following washout of gabazine at DIV14, numerous topological characteristics are recovered. Within our simulations, the washout effect is reflected as a shift towards a more canonical lognormal connectivity distribution^103^.

### Probing generative wiring principles across different human neuronal networks

As shown in the previous section, patterns of neuronal spiking dynamics may have an effect on how rodent neurons form functional connectivity *in vitro*. However, to what extent this idea translates to, for example, networks comprising specific kinds of human neurons and their respective/varying spiking dynamics^105,106^ remains unclear. We start to address this question by applying GNMs to purified human iPSC-derived neuron/astrocyte co-cultures. GNM analyses were performed at a time point, at which such cultures reach a state of relative maturity (DIV 28)^65^. The dataset for this analysis comprises purified glutamatergic neurons (GNs, n=8), motor neurons (MNs, n=7), and dopaminergic neuronal cultures (DNs, n=6). We also included slice cultures derived from 4-month-old human embryonic stem cell-derived cerebral organoids (hCOs, n=6 slices). Previous studies have indicated that hCOs develop functional networks with increasing complexity from as early as 90 days *in vitro*^107,108^. **Figure 7a** shows an immunohistochemical staining of a DIV 21 human iPSC-derived DN culture, expressing neuronal and astrocytic markers (MAP2, GFAP, and TH); **Figure 7b** shows stainings for hCOs slices (Tau, NeuN, and GFAP).

**Figure 7.**
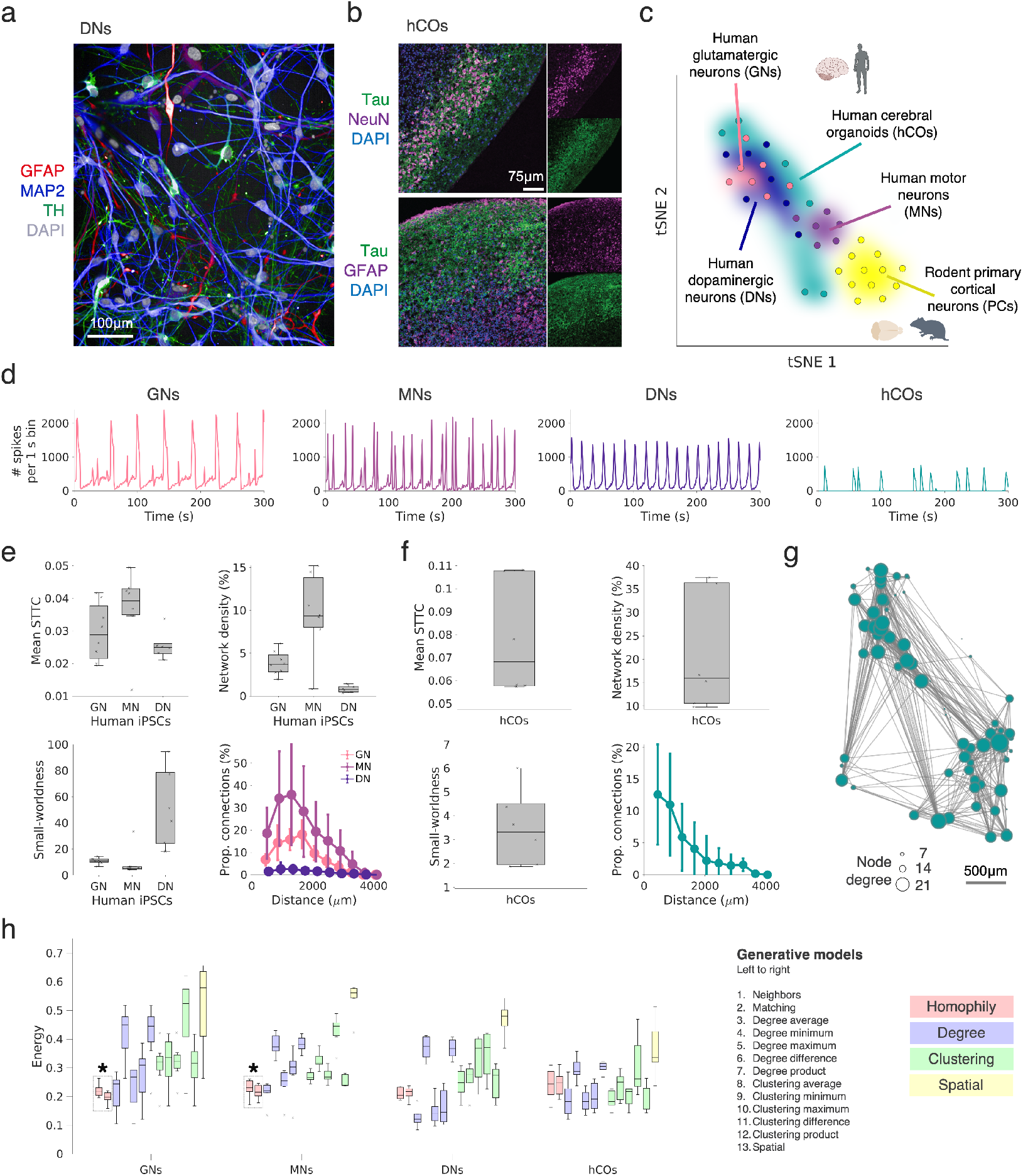
Generative network modeling across human neuronal cultures and cerebral organoids. **a** An immunohistochemical staining of a single hCO slice (Tau in green, NeuN in purple and DAPI in blue; bottom panel: Tau in green, GFAP in purple and DAPI in blue). **b** Clustering of human and rodent neuronal cultures based on a t-distributed stochastic neighborhood embedding (tSNE). Each dot corresponds to one culture; similarity was estimated by correlating the spike train autocorrelograms across all datasets and by then applying the tSNE. Rodent PC neuronal networks (yellow, dense platings, n=12, DIV 28), human iPSC-derived dopaminergic (dark-blue, n=6), motor (purple, n=7) and glutamatergic neurons (pink, n=8) form clear clusters in 2D tSNE space; human ESC-derived hCOs (green, n=6) were more scattered. **c** Immunohistochemical staining of a control dopaminergic neuron (DN) network (top panel: expression of GFAP in red, MAP2 in blue, TH in green, and DAPI in gray). **d** Representative population activity plots for networks of glutamatergic (left), motor (left middle), dopaminergic (right middle) iPSC-derived neuronal cultures and a hCO slice (right). **e** Global topological measures of human iPSC-derived neuronal networks at DIV 28: the mean STTC (top left), network density (top right), small-worldness (bottom left) and proportion of extant connection by distance (bottom right). **f** STTC functional connectivity measures and small-worldness of hCO slices including: average STTC (top left), network density (top right), small-worldness (bottom left), and connectivity as a function of distance (bottom right). **g** Functional connectivity graph inferred from a 120-day-old hCO slice. **h** Fits across generative model rules in human iPSC-derived neuronal networks (DIV 28; n=6 for each cell line, left, middle left and middle right) and hCOs (n=6, right). The energy of the top n=1 performing simulated networks are shown. Boxplots represent the median and IQR. Outliers are demarcated as small black crosses, and are those which exceed 1.5x the IQR away from the top or bottom of the box. The asterisk reflects homophilic rules which demonstrate statistically lower energy (p<0.05) than other rules. All statistics are provided in **Supplementary Table 7**.

**Figure 7c** provides an overview of the human and rodent neuronal spiking dynamics. Following a t-distributed stochastic neighbor embedding (tSNE) analysis, we find that networks can be clustered well according to their spiking dynamics. The tSNE analysis is based on spike-train autocorrelograms, derived from the aggregated activity of each neuronal network^109^ (see **Methods; *Autocorrelogram analysis***). Example network activity plots, illustrating the different firing dynamics and corresponding autocorrelograms, are shown in **Figure 7d** and **Supplementary Figure 15a**. Overall, activity-based tSNE clustering allows grouping of different monolayer human neuronal cell lines in space; greater spread/variability is observed for the hCO slices. MN dynamics appear closest to rodent primary cortical neurons (PCs) in the tSNE space, which this is reflected by their similarity in ongoing spiking dynamics (**Supplementary Figure 15b**). Representative examples of the observed differences in population activities across different human neuron cultures are depicted in **Figure 7d**.

As for the rodent cultures, we constructed STTC functional connectivity graphs for all human neuron cultures, and assessed key connectivity metrics and topology. **Figure 7e** and **Figure 7f** show these network statistics respectively; **Figure 7g** provides an example of a single constructed hCO functional network. It is notable that human iPSC-derived neuronal networks did not differ significantly in average STTC (ANOVA, p=0.0912). However, we find a significant difference in network density (p=1.02×10^-4^) and topological metrics, such as the small-world index (p=1.51×10^-4^). MNs show the highest network density, as inferred by the STTC, and small-worldness values within the range previously observed for the rodent PC networks.

**Figure 7h** depicts the GNM results across all human monolayer and organoid cultures. Overall, generative model findings approximately mirror the similarity in their underlying spiking dynamics (**Figure 7d**), in that human MNs show an energy profile across generative models aligning with rodent cultures, followed by GNs (**Figure 7h**, left and middle-left; homophilic rules p<3.70×10^-3^ for all pairwise comparisons and Cohen’s *d*>0.772 reflecting large effect sizes). In contrast, DNs and hCOs, which show differing underlying dynamics, exhibit a very distinct profile with the degree average method achieving the lowest energy, but no non-spatial class of generative rules providing the statistically lowest energy (**Figure 7h**, middle-right and right). As noted before, hCOs exhibited significant variability in their spiking dynamics and topology, and provide an energy profile somewhat resembling randomly rewired graphs (**Supplementary Figure 5b**, right). All statistical findings are provided in full in **Supplementary Table 7**.

In summary, GNMs based on homophilic wiring mechanisms best recapitulate functional network connectivity *in vitro* in rodent PC networks, particularly, at later developmental stages (DIV 14 onwards). In few cases, including some immature rodent PC networks and human cultures, the degree-average-based model performs well, if not best. This may reflect differences in the underlying spiking dynamics (**Supplementary Figure 15**). Human GN and MN networks show the closest resemblance to rodent PC networks, and as such, mirror their underlying homophilic generative mechanisms. Results in hCOs are as yet inconclusive, likely due to their observed variability in spiking activity and immature functional connectivity.

## DISCUSSION

In the current study we applied large-scale electrophysiological recordings to track and characterize single-unit functional connectivity as neuronal networks develop *in vitro*. We systematically tested which candidate topological attachment mechanisms could explain this developing self-organization, using a range of generative network models to simulate network formation *in silico*. In the majority of cases tested, we found that a model utilizing an homophilic attachment mechanism^45^ performed best. This model accurately captures the developmental trajectories of neuronal networks, their local topological organization and highlights how neuronal variability is likely critical to the emergence of the canonical lognormal distribution^103^ of neuronal functional connectivity within networks. The apparent symmetry between these findings and previous work at various scales of analysis^51,53,60,61,63^ and species^54,110^ may have implications for the study of brain development.

### Topological self-similarity as a driver of in vitro topology

In line with previous work, we find that functional connectivity increased with development, and that developing neuronal networks *in vitro* exhibited typical characteristics of complex network architecture seen across numerous scales^66,111^. Our work demonstrates that, beyond the other tested models, homophily best recapitulates complex functional network topology *in vitro*.

Interestingly, we find that the degree-average model tends to consistently follow the performance of homophily in cultures that are relatively less complex (e.g., immature rodent cultures and human single-cell-type cultures). Conceptually, the degree-average generative model prefers connection formation when both neurons simultaneously have large numbers of connections (given that γ>0). Importantly, the degree-average model (as with all other models apart from homophily) treats the underlying computation for how networks form connections as entirely independent. That is, the computation for the probability to wire is made not with direct respect to any other node pair—it is made on each neuron before then performing some other operation (e.g., taking the average, or maximum). In contrast, homophily is a function made with respect to the *direct* relationship between the connections of the neurons. This is a subtle but important theoretical distinction, because it highlights, how homophily can occur via the local communication between neurons, where signals are propagated via their connections. This is likely critical, as any generative mechanism by which complex neurobiological networks develop are likely to emerge from these interactions between the local components over time^26^—without a central mechanism aiming to optimize its global network properties^112^.

A related observation is that the homophily heuristic—much like in social networks^113,114^— enables each part of the network to interact with its local environment without requiring inordinate computational resources. Indeed, homophily has been shown to provide an efficient trade-off capable of producing small-world networks through which information can propagate efficiently^91,92^. Under this view, as limits to *local-knowledge* and *computational capacity* hold for any interacting developing system, homophily becomes a generative heuristic for any sufficiently large network. Notably, this resonates with accounts of Hebbian learning^115,116^ and spike-timing dependent plasticity (STDP)^117^ whereby neurons wire with each other as a function of *similarity^118^* to themselves (e.g., concurrent or temporally precedent neuronal firing, respectively) provided that neurons are sufficiently close in space.

### Models of developing networks across scales, species and time

The combination of generative modeling and graph theory allows us to use *in silico* simulations as a lingua franca to probe micro-connectomic self-organization^13^. Comparative studies have examined economic accounts of connectomic organization across different species^20^—such as in the worm *C. elegans*^37,91,119^, larval zebrafish^29^, mouse^120^, macaque^54^ and human connectome^51–53,56,58–61,63^. For example, Nicosea, *et al*.^91^ modeled the growth of *C. elegans* using the known birth times of its somatic neurons—finding that as the body of the animal progressively elongates, the cost of longer-distance connections become increasingly penalized. In human brain scans, Oldham, *et al*.^60^ incorporated known early changes in brain macroscopic geometry and other physiological measures of homophily (e.g., correlated gene expression) to improve an additive generative model’s network embedding (also see^52^). These works have highlighted the benefit of incorporating specific developmental changes, that are specific to the organism, within a growth model able to simulate developmental outcomes.

### Homogeneous spiking dynamics may lead to stochastic wiring through reduced parameter magnitude

In line with previous findings, the inhibition of GABA_A_ receptors led to increased neuronal synchronization compared to untreated cultures^97,121^. Despite this difference in population dynamics the homophily generative model better reproduced network topology than the other models, albeit with less error in the control networks. Crucially, GABA_A_ receptor block did significantly push the model parameter, η, closer to zero. Reduced η magnitude indicates a weaker preference for short-distance connections in gabazine-treated networks compared to controls, whilst still preferring shorter connections as indicated by η being negative. One explanation might be that increased synchronicity with gabazine application leads to functionally less distinguishable spike dynamics between neurons—hence the connection probability distribution was flatter and shifted to the right relative to the probability distribution of controls. One might expect that without outstanding high-probability connections to make, connection formation will be driven by cost with neuronal units preferentially connecting with more proximal units with long-range connections having little added value. However, η magnitude decreased rather than increased with gabazine treatment. This indicates that gabazine-treated cultures showed a weaker rather than stronger preference for short-distance connections compared to controls. One explanation could be that gabazine-treated cultures fail to deselect less homophilic longdistance connections. Consistent with this, gabazine-treated cultures showed a longer total edge length, reflecting more long-distance, high-cost connectivity than controls. Together, these results support that GABA_A_ receptor-mediated activity influenced connection specification that can elicit the same network topology more efficiently by maximizing value relative to cost of connections.

Interestingly, previous GNM work at the whole-brain scale has shown that lower magnitude wiring parameters are associated with poorer cognitive scores^51^, age^51,53^ and a diagnosis of Schizophrenia^61,63^. Perhaps, inhibitory hub neurons fail to implement the principle of homophily in the same way, thus constraining the emergence of small-worldness which has also been related cognitive function^122,123^. This may suggest convergent evidence for how developmental randomness, intrinsic to how developing parts interact with each other, may influence functional outcomes^124^. A remaining challenge in the field is to be able to directly parse the extent to which stochasticity and specific economic trade-offs may influence network outcomes under different conditions.

### Limitations and future work

There is a lack of consensus as to how functional connectivity can be inferred from the spontaneous activity of neurons developing *in vitro*^125^. In the present study, we utilized the spike time tiling coefficient (STTC^73^) which was developed to improve on some of the limitations of traditional metrics for the coupling between neurons, such as the correlation index^126^. Crucially, our goal was not to infer synaptic connections, direct structural connections nor reconstruct the underlying circuit of synaptic (and autaptic^127^) connections present. Rather we sought to infer relationships in temporal patterns of spiking activity between neuronal units that may be relevant to network emergence and its functioning. Thus, the present analysis sits at a larger spatial scale than the synaptic or circuit level within the spatial and temporal structure of the brain. Nevertheless, in **Supplementary Figure 5**, we show that our results can be replicated with transfer entropy-based methods^75,128^, i.e., measures of effective connectivity, which may better correspond to aggregated connections weights between neurons. Furthermore, similar results were found in standard density MEAs, though neuronal units were not tracked over time^67^.

An important outstanding question is how the role of GABA_A_ receptor-mediated activity changes with development, and whether there is any link to the observed findings. From DIV 7 to DIV 14, homophily became more distinguishable from other rules in terms of its recapitulation of network topology in rodent PC networks. This coincides with the gradually increasing proportion of GABAergic synapse switching from depolarization to hyperpolarization during this time in rodent cultures^100,129^. However, we inhibited GABA_A_ receptors chronically from DIV1 and found homophilic rules still provided the best fit. Therefore, whilst GABA_A_ receptor-mediated inhibition may play a role in de-coupling neurons and hub node function^93,94,97^, further work is required to understand the precise relationship between inhibition mediated by different receptor subtypes (e.g. GABA_B_ receptor-mediated), cell types and how this changes wiring parameter magnitude, and ultimately network topology, over development.

Another potential limitation relates to the application of our modeling approach to human monolayer and organoid cultures. The human iPSC-derived neuronal networks consist of purified cell-types, which is clearly artificial. As reflected by the lack of connectivity and topology in some of these cultures, such as the DN networks, which also did not show homophilic wiring, there is perhaps little surprise they performed differently. A more veridical account will likely come from cultures with mixed iPSC lines exhibiting more complex spiking dynamics arising from defined cell-type driven heterogeneity^130^. It is also possible that the spatial extent of the recording from 3D hCOs onto the 2D HD-MEA system may have limited our network inferences.

Despite these caveats, our present work shows that homophilic generative models *per se* are appropriate growth models for *in vitro* neuronal networks as they were capable of recapitulating key statistical properties—both at the local and global level. However, as noted in prior GNM studies^51,60^, a significant future advance in this research area will come from weighted generative network models capable of recapitulating weighted topological architectures. Such an approach would allow for both the tuning of connection weights over developmental time—a clear principle of network maturation^119^—but also enable further study of how developing network topology, genetics^52,60,131^ and information processing^54,132^ together explain neuronal network organization across scales.

In conclusion, we find that the complex topology of developing rodent and many human neuronal networks *in vitro* can be simulated via a simple homophily generative model, where neurons aim to maximize locally shared connectivity within an economic context. With this, and prior research at the macroscopic level in mind, we suggest that homophily wiring rules provide a compelling isomorphic explanation for any decentralized, locally computing, developing system.

## METHODS

### High-density microelectrode arrays

Two types of CMOS-based high-density microelectrode array (HD-MEA) recording systems, produced by MaxWell Biosystems (Zurich, Switzerland), were used in the present study^64,133^. The single-well HD-MEA MaxOne, consisting of 26,400 low-noise electrodes with a center-to-center electrode pitch of 17.5 μm, arranged in a 120 x 220 electrode array structure. This HD-MEA can record simultaneously from a total of 1024 (user-selected; 3.85 × 2.10 mm^2^ sensing area) readout-channels at 20 kHz; for more technical details see previous studies^64,133^. The second recording system was the multi-well HD-MEA MaxTwo (MaxWell Biosystems, Zurich, Switzerland), comprising the same number of electrodes and readout-channels and electrode specifications as MaxOne for each well. With this system it is possible to simultaneously record from six wells at a time and at a sampling rate of 10 kHz. To decrease the impedance and to improve the signal-to-noise ratio (SNR), electrodes were coated with platinum black^64^.

### Rodent primary cortical neuronal cultures

Before plating, HD-MEAs were sterilized in 70% ethanol for 30 minutes and rinsed three times with sterile water. To enhance cell adhesion, the electrode area of all HD-MEAs was treated with poly-D-lysine (PDL, 20 μL, 0.1 mg mL^-1^; A3890401, Gibco, ThermoFisher Scientific, Waltham, USA) for 1 hour at room temperature and then rinsed three times with sterile water. Next, 10 μL Geltrex (A1569601, Gibco, 0.16 mg mL^-1^) was pipetted on each array and again left for about one hour at room temperature. For the main analysis of the paper, we used rodent primary cortical (PC) neurons prepared as previously described^64^. Briefly, cortices of embryonic day (E) 18/19 Wistar rats were dissociated in trypsin with 0.25% EDTA (Gibco), washed after 20 min of digestion in plating medium (see below), and triturated. Following cell counting with a hemocytometer, either 50,000 cells (sparse plating condition) or 100,000 cells were seeded on each array, and afterwards placed in a cell culture incubator for 30 min at 37°C/5% CO_2_. Next, plating medium was added carefully to each well. The plating medium contained 450 mL Neurobasal (Invitrogen, Carlsbad, CA, United States), 50 mL horse serum (HyClone, Thermo Fisher Scientific), 1.25 mL Glutamax (Invitrogen), and 10 mL B-27 (Invitrogen). After two days, half of the plating medium was exchanged with growth medium containing 450 mL D-MEM (Invitrogen), 50 mL horse serum (HyClone), 1.25 mL Glutamax (Invitrogen) and 5 mL sodium pyruvate (Invitrogen). Across all experiments, the medium was then exchanged twice a week, at least one day before the recording sessions. All animal experiments were approved by the veterinary office of the Kanton Basel-Stadt, and carried out according to Swiss federal laws on animal welfare. A summary of the data used is provided in **Supplementary Table 1.**

### Human induced pluripotent stem cell-derived neuronal cultures

Three different human iPSC-derived neuronal cell lines were included in the study: iCell DopaNeurons, iCell Motor Neurons and iCell GlutaNeurons, all commercially available from FUJIFILM Cellular Dynamics International (FCDI, Madison, USA). All neural cells were cocultured with human iCell Astrocytes (FCDI, see above). Cell plating: Cell plating medium consisted of 95 mL of BrainPhys Neuronal Medium (STEMCELL Technologies, Vancouver, Canada), 2 mL of iCell Neuronal Supplement B (FCDI), 1 mL iCell Nervous System Supplement (FCDI), 1 mL N-2 Supplement (100X, Gibco), 0.1 mL laminin (1 mg/mL, Sigma-Aldrich) and 1 mL Penicillin-Streptomycin (100X, Gibco). Neurons and astrocytes were thawed in a 37°C water bath for 3 minutes. The cells were then transferred to 50 mL centrifuge tubes, and 8 mL plating medium (at room temperature) was carefully added. Cell suspensions were centrifuged at 380 x g (1600 RPM) for 5 minutes, and the supernatant was aspirated. Cell pellets were then resuspended in plating medium and combined to achieve a final concentration of 10,000 neurons and 2,000 astrocytes per μL. Finally, 100,000 neurons and 20,000 astrocytes were seeded per HD-MEA by adding 10 μL of the prepared solution, after removing the Geltrex droplet. After incubating the cells for one hour at 37°C/5% CO2, another 0.6 mL (small well MaxOne) / 1.2 mL (large well MaxOne) of plating medium was added. Half of the medium was changed twice a week.

### Human cerebral organoid slice cultures

Human embryonic stem cell (hESC)-derived cerebral organoids (hCOs) were generated from a commercially available hESC stem cell line (Takara Bio, Osaka, Japan), using the STEMdiff cerebral organoid kit (STEMCELL Technologies) following the manufacturer’s instructions. Slices were obtained from 120-day old hCOs. Single organoids were first transferred from maturation medium to ice-cold BrainPhys (STEMCELL Technologies) using cut 1000 μl pipette tips. Next, cross-sectional 500-μm-thick slices were cut from hCOs using a sterile razor blade and collected in petri dishes filled with BrainPhys medium at room temperature. Before the plating, HD-MEAs were sterilized in 70% ethanol for 30 minutes and rinsed 3 times with distilled water. To improve tissue adhesion, arrays were coated with 0.05% (v/v) poly(ethyleneimine) (Sigma-Aldrich) in borate buffer (pH 8.5, Thermo Fisher Scientific) for 30 minutes at room temperature, rinsed with distilled water, and left to dry. To attach hCOs on HD-MEAs, we applied a thin layer of Matrigel (Corning) to the center of the HD-MEA and then transferred individual organoid slices to the coated HD-MEAs. After positioning the tissue, we placed a tissue “harp” on top of the organoid slice and applied several drops of recording medium (STEMCELL Technologies, #05793) around the organoid. HD-MEAs were then covered with a lid and placed in a humidified incubator at 37°C, 5% CO2/95% air for 30 minutes, before adding more medium to a final volume of 2 ml per chip. Half of the recording medium was changed every 2-3 days.

### Immunohistochemistry

Rodent PC neurons were stained as previously described^64^. Briefly, PC neurons were fixed using 4% paraformaldehyde solution (ThermoFisher, #FB001). Samples were permeabilized and blocked using a PBS 10X (ThermoFisher, #AM9625) solution containing 10% normal donkey serum (NDS) (Jackson ImmunoResearch, West Grove, USA, #017000001), 1% bovine serum albumin (BSA) (Sigma-Aldrich, 0 5482), 0.02% Na-Az (Sigma-Aldrich, #S2002) and 0.5% Triton X (Sigma-Aldrich, #93443). Permeabilization facilitated antigen access to the cell, while blocking prevented non-specific binding of antibodies to neurons. Primary and secondary antibodies were diluted in a PBS solution containing 3% NDS, 1% BSA, 0.02% Na-Az and 0.5% Triton X. The used antibodies are also listed in **Supplemental Table 8**. Note, immunohistochemistry was performed on control PC cultures prepared as previously outlined^65^.

Human iPSC-derived neurons were fixed using 8% PFA solution (#15714S, Electron Microscopy Sciences) and blocked for 1 hour at room temperature (RT) in blocking buffer containing 10% normal donkey serum (NDS) (Jackson ImmunoResearch, West Grove, USA, #017-000-001), 1% bovine serum albumin (BSA) (#05482, Sigma-Aldrich), and 0.2% Triton X (Sigma-Aldrich, #93443) in PBS (ThermoFisher Scientific, #AM9625). Primary antibodies (**Supplementary Table 8**) were diluted in a blocking buffer and incubated overnight at 4°C. Samples were washed three times with 1% BSA in PBS and incubated with the secondary antibody (**Supplementary Table 8**) diluted in blocking buffer for 1 hour at RT. After three additional washes with PBS, DAPI was added for 2 min at RT (1:10000). Images were acquired using the Opera Phenix Plus High-Content Screening System (cat. HH14001000, PerkinElmer, Waltham, MA, USA).

hCOs were fixed using 4% paraformaldehyde (PFA) for 4 hours at room temperature, washed with PBS and immersed in 30% sucrose solution at 4 °C overnight. PFA-fixed organoids were embedded in OCT compound (Sakura Finetek, Alphen aan den Rijn, Netherlands, #4583) and stored at −80 °C. 10 μm sections were cut on a cryostat and collected on Superfrost plus slides (Thermo Scientific, #22-037-246). For immunohistochemistry, sections were permeabilized in 0.1% Triton X-100 and blocked with animal-free blocker (Vector Laboratories, Burlingame, CA, USA, #SP-5030-250). Slides were incubated with primary antibodies for 1 hour at room temperature. Sections were washed in PBS and further incubated with secondary antibodies for 1 hour at room temperature. After washing with PBS, sections were incubated with PureBlu DAPI (Bio-Rad, Hercules, CA, USA, #1351303) for 3 minutes and mounted with ProLong Gold antifade mounting medium (Thermo Scientific, #P36930). Fluorescence images were acquired with a SP8 confocal microscope (Leica, Wetzlar, Germany). The primary and secondary antibodies used for hCO stainings are listed in **Supplementary Table 8**.

### Scanning electron microscope imaging

Fresh tissue samples were fixed in 2.5% glutaraldehyde solution (Sigma-Aldrich, St. Louis, USA) overnight. After fixation, the samples were dehydrated in ascending acetone series (50%, 70%, 80%, 90%, 95%, 100%), and critically point dried (CPD; Quorum Technologies, West Sussex, UK), using CO2 as the substitution fluid. The procedure is generally suited for SEM preparation and ensures that surface structures of animal tissue samples are preserved in their natural state, i.e., without shrinkage, distortion or dissolution. After CPD, specimens were carefully mounted on aluminum stubs using double sticky carbon-coated tabs as adhesive (Plano, Wetzlar, Germany). Thereafter, they were coated with gold-palladium in a sputter device for 45 seconds (Bio-Rad SC 510, Munich, Germany). SEM analyses were carried out with a Zeiss Digital Scanning Electron Microscope (SUPRA 40 VP, Oberkochen, Germany) in SE2 mode at 5-10 kV.

### Electrophysiological recordings

In order to track the development of functional connectivity of *in vitro* neuronal networks on HD-MEAs, we performed weekly recordings, starting one week after plating. In order to select a network recording configuration, we performed whole-array activity scans, i.e., series of 1-minute long high-density recordings, covering all 26,400 electrodes of the HD-MEA, using the MaxLab Live software (MaxWell Biosystems). To select recording electrodes, we estimated the multi-unit activity for each electrode using an online sliding window threshold-crossing spike-detection algorithm (window length: 1024 samples; detection threshold: 4.5 × the root mean squared error (RMSE) of the noise of the 300-3000 Hz bandpass filtered signal). After the activity scan, we selected up to 1024 readout-electrodes, based on the detected average activity and a ranking of the inferred, average amplitude values. Additional high-density network recordings, consisting of 4 x 4 electrode blocks (17.5 μm pitch), were acquired for the tracking experiments (see below). The duration of the HD-MEA network recordings was about 30 minutes; an overview on the different datasets is provided in **Supplemental Table 1**. The PC neuronal network and the hCO data were acquired by MaxTwo multi-well plates (MaxWell Biosystems); the human iPSC-derived neurons (glutamatergic, motor and dopaminergic neurons) were recorded on single-well MaxOne HD-MEAs (MaxWell Biosystems).

### Pharmacological experiments

Pharmacological experiments with the GABA_A_ receptor blocker gabazine (SR 95531 hydrobromide, Sigma-Aldrich, #104104509), were performed on sparse (50,000 per well) primary cortical (PC) neuronal cultures. Three cultures were treated with 1 μM gabazine one day after plating and tracked until DIV14; media+gabazine exchanges were performed 2-3 times per week.

### Spike-sorting and post-processing

All HD-MEA network recordings underwent an initial quality control to assess the overall noise level and signal stability of each recording. Next, we used the software package Kilosort 2 (KS2)^72^ to spike sort data, applying default parameters. For the developmental tracking analyses, we concatenated all recordings (i.e., DIV7, 10, 12, and 14 for the PC cultures at 50k plating density, and DIV14, 21 and 28 for the PC cultures plated at 100k per well). After spike sorting, we inferred array-wide spike-triggered averages (STAs) for all units labeled as ‘good’ by KS2. Next, we calculated the spatial similarity between all detected units/STAs to minimize the influence of potential cluster splits that might have occurred during spike sorting of bursty spontaneous activity. The spatial similarity among the inferred array-wide templates was probed by the normalized pairwise maximum cross-correlation: units/STAs that showed a similarity *r* >0.75 and had at least 5 electrodes in common underwent an iterative elimination process using a simple clustering heuristic^134^. Please see **Supplementary Table 1** for a summary of the data sets used in this study, and **Supplemental Figure 1** for the number of trackable units for both datasets.

### Firing rate and burst statistics

Firing rates across each neuronal unit were calculated as the total number of spikes per unit time (in seconds) in the entire recording. Array values were calculated as the mean across all active units (firing rates >0.01 Hz). Burst rates were calculated using a maximum interspike interval (ISI) method^135^ based on the ISI between every Nth spike (ISI_N_)^136^. The ISI_N_ threshold for determining the onset/offset of bursting activity was determined by finding the local trough in the bimodal logISI distribution (see **Supplemental Figure 11b**). The two peaks, at short ISIs and long ISIs represent more high frequency bursting and regular activity, respectively. The coefficient of variation (CV) of interburst intervals (IBIs) was calculated as the standard deviation of IBIs relative to the mean IBI in a given neuronal unit; the array value was the mean of this across all neuronal units.

### Functional connectivity inference

To detect pairwise correlations in spike trains, here referred to as functional connectivity, we computed the spiketime tiling coefficient (STTC)^73^. The STTC aims to mitigate potential confounding in basic correlation indices introduced by different firing rates, by quantifying the proportion of spikes in one train which fall within ±Δt (the synchronicity window) of another. It is given by:

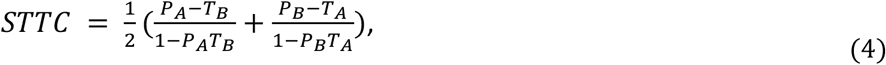

where T_A_ is the proportion of total recording time which lies within ±Δt of any spike from A (T_B_ is calculated similarly). P_A_ is the proportion of spikes from A which lies within ±Δt of any spike from B (P_B_ is calculated similarly). The synchronicity window, Δt, is the only free parameter in the STTC calculation. In the present study, we used a Δt=10 ms. A visualization of the STTC calculation is provided in **Figure 1h**; STTC was calculated using publicly available Matlab code^82^. We used permutation-based testing to determine the significance of connections. For a given neuronal unit’s spike train, spike times were randomly jittered by ±10ms to create a surrogate spike train, using code provided by the Neural Complexity and Criticality Toolbox^137^. This was repeated for each neuronal unit for 1000 permutations. To calculate significance of pairwise functional connectivity, experimentally inferred STTC values were compared to the distribution of surrogate SSTC values. A significance value of p < 0.01 was used as a cutoff to binarize functional connectivity matrices and calculate network related analysis throughout the manuscript; only units with firing rates >0.01 Hz were considered.

### Network statistics

In **Figure 2a** we provide a visualization of key graph theoretical metrics relevant for this study. Here we provide both a written and mathematical definition for each measure used. Each statistic was calculated using the Brain Connectivity Toolbox^78^:

#### Degree

The degree is the number of edges connected to a node. The degree of node *i* is given by:

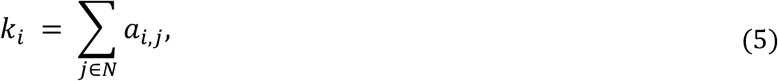

where *α_i,j_* is the connection status between *i* and *j*. *a_i,j_* =1 when link *i,j* exists (when *i* and *j* are neighbors); *α_i,j_* = 0 otherwise (*α_i,i_* = 0 for all *i*).

#### Clustering coefficient

The clustering coefficient is the fraction of a node’s neighbors that are neighbors of each other. The clustering coefficient for node *i* is given by:

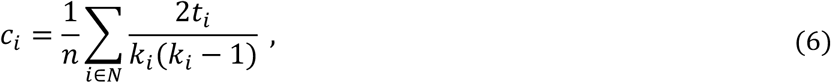

where *C_i_* is the clustering coefficient of node *i* (*C_i_* = 0 for *k_i_* < 2).

#### Betweenness centrality

The betweenness centrality is the fraction of all shortest paths in the network that contain a given node. Nodes with high values of betweenness centrality therefore participate in a large number of shortest paths. The betweenness centrality for node *i* is given by:

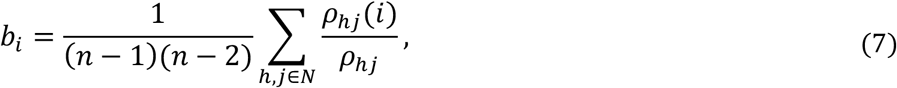

where *ρ_hj_* is the number of shortest paths between *h* and *j*, and *p_hj_*(*i*) is the number of shortest paths between *h* and *j* that pass through *i*.

#### Edge length

The edge length is the total edge lengths connected to a node. It is given by:

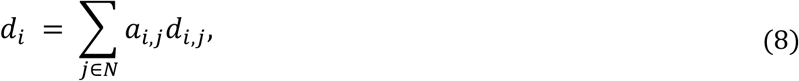

where *d_i,j_* is the Euclidean distance between *i* and *j*. The Euclidean distances of functional connectivity graphs inferred in the present study are depicted in **Supplementary Figure 3**.

#### Global efficiency

The global efficiency is the average of inverse shortest path length. It is given by:

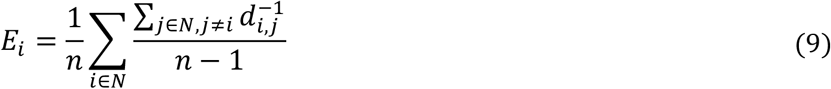

#### Matching

The matching index computes the proportion of overlap in the connectivity between two nodes. It is given by:

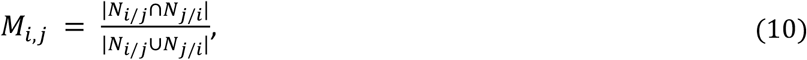

where *N_i/j_* refers to neighbors of the node *i* excluding node *j*. Where global measures of matching have been used, we averaged across the upper triangle of the computed matching matrix.

#### Small-worldness

Small-worldness refers to a graph property where most nodes are not neighbors of one another, but the neighbors of nodes are likely to be neighbors of each other. This means that most nodes can be reached from every other node in a small number of steps. It is given by:

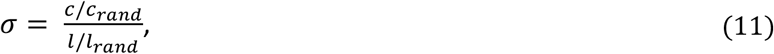

where *c* and *c_rand_* are the clustering coefficients, and *l* and *l_rand_* are the characteristic path lengths of the respective tested network and a random network with the same size and density of the empirical network. Networks are generally considered as small-world networks at σ>1. In our work, we computed the random network as the mean statistic across a distribution of n=1000 random networks. The characteristic path length is given by:

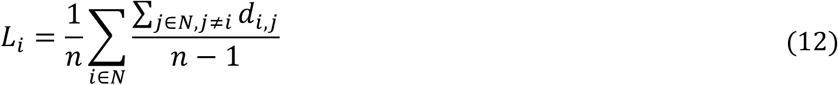

#### Modularity

The modularity statistic, Q, quantifies the extent to which the network can be subdivided into clearly delineated groups:

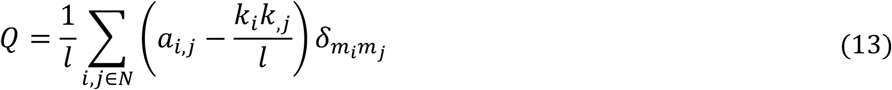

where *m_i_* is the module containing node *i*, and *δ_m_i_m_j__* = 1 if *m_i_* = *m_j_*, and 0 otherwise.

#### Participation coefficient

The participation coefficient is a measure of diversity of intermodular connections of individual nodes, where community allocation was determined via a Louvain algorithm, with a resolution parameter γ = 1, which aims to form a subdivision of the network which maximizes the number of within-group edges and minimizes between group edges.

### Generative network modeling

The generative network model can be expressed as a simple wiring equation^51,53,56,61^, where wiring probabilities are computed iteratively by trading-off the cost of forming a connection, against the value of the connection being formed in terms of a network topology term. Connections are added iteratively according to these wiring probabilities. It is given by the wiring equation as provided in **Equation 1**. The *D_i,j_* term represents the “costs” incurred between neurons modeled as the Euclidean distance between tracked units (**Supplementary Figure 3**). The *K_i,j_* term represents how neurons “value” each other, given by an arbitrary topological relationship which is postulated *a priori* (also termed, “wiring rule” given mathematically in **Supplementary Table 2**). *P_i,j_* reflects the probability of forming a fixed binary connection at the current time step. The simulation continues until the simulated network has the same number of connections of the observed network. The *D_i,j_* term remains constant during the simulation while the *K_i,j_* term updates at each time point (and therefore also the *P_i,j_* term).

### Cost functions

In the present study, we make a distinction between simulated networks which mirror the statistical distributions of observed networks and those which mirror the topological organization of those statistics. The former can be accessed via a previously used energy equation^53^ whereby the model fit is given by the “worst” of the four *KS* distances assessed, given by **Equation 2**. *KS* is the Kolmogorov-Smirnov statistic between the observed and simulated networks at the particular η and γ combination used to construct the simulation, defined by the network degree *k*, clustering coefficient *c*, betweenness centrality *b* and Euclidean edge length *d*. Notably, the KS distance between two vectors simply considers their statistical *distributions*.

In **Supplementary Figure 6**, we further assess the ability of the best performing generative models in each class (spatial, matching, clustering average and degree average) to recapitulate network statistics as included in the energy equation, but also two measures outside (local efficiency and participation coefficient). We did this via a Monte Carlo sampling procedure^83^. First, we took the top n=99 performing simulations for each sparse rodent culture’s model considered, and computed each of the six local statistics as shown in **Supplementary Figure 6** as cumulative density plots. For each statistic, we computed a KS statistic between the observed local statistics distribution and an average of the statistics of the 99 simulations. We then undertook 99 individual leave-one-out iterations in which we replaced a single simulation of the 99 with the observed distribution. For each of the 99 permutations, we computed the same statistic, forming a null distribution. We then calculated a p_rank_ by ranking how close the original observed statistic was to the mean of this computed null distribution (i.e., how close was the observation to the middle of the null). This was computed for each culture and statistic, for each of the considered generative models. We then quoted the median p_rank_ across cultures.

Later in the study, we provide an alternative but simple cost function which does not assess distributions of statistics, but instead assesses the *topological fingerprint dissimilarity* of these network statistics. The *topological fingerprint (TF*) matrix is calculated as a Pearson’s correlation matrix between each pair-wise combination of the local statistics. In our study, we used six common network statistics to form this correlation matrix, however, in principle, these can be extended to any number or range of local statistical measures. The construction of the *TF* is visualized in **Figure 5a**. The *TF_dissimilarity_* is then calculated as the Euclidean norm^92^ of the difference between the observed and simulated *TF* matrices. This is given in **Equation 3**.

### Parameter selection

We optimized η and γ using a Voronoi tessellation procedure as used in prior work^53^. This procedure works by first randomly sampling the parameter space and evaluating the model fits of the resulting simulated networks, via the energy equation. As there is little prior literature that can be used to guide the present study, we considered a wider range of parameter values, with η values in the range from −10 to 10 and γ values in the range −10 to 10. Following an initial search of 4000 parameters in this space, we performed a Voronoi tessellation, which establishes two-dimensional cells of the space. We then preferentially sampled from cells with better model fits according to the energy equation (see^53^ for further detail). Preference was computed with a severity of α = 2 which determines the extent to which cell performance led to preferential sampling in the next step. This procedure was repeated a further four times, leading to a total of 20,000 simulations being run for each considered network across the 13 generative rules as described in **Supplementary Table 2**.

### Generative probability distributions

In **Figure 6d**, we show the mean probability score (*P_i,j_*) distributions within the generative models fit to gabazine and control networks. This was calculated by measuring the *P_i,j_* across all node pairs *i* and *j* in the network, in 1% intervals, before plotting the average distribution of *P_i,j_* across these timesteps. In **Supplementary Figure 13e,f**, we show each distribution of these probability distributions (that was averaged to provide comparisons in **Figure 6d**). Note that the probability score distribution flattening means there are more edges with higher probabilities of being connected, leading to decreased specificity of future wiring. This flattening effect is equivalent to the network outcomes being more random.

### Autocorrelogram analysis

Autocorrelogram analysis was carried out using the CellExplorer Matlab Toolbox CGG function^109,138^. First, spike times were concatenated cumulatively across units to give a single vector of spike times. Spikes were summed into consecutive one millisecond bins giving a vector where each element is a one millisecond bin containing the number of spikes occurring in the network at each time point. This vector, *v*, was then correlated with itself plus a lag value, *x*. The range of lag values tested was −500 – 500 ms. Lag values between −1 – 1 ms were removed to impose a refractory period—hence, these values are 0 in ***Supplementary Figure 15***. For example, where *x* is 5 ms, the CCG function is the sum of *v_i_* and *v_t+x_* across all time points, *t*. This gives a vector of CCG values, corresponding to spatiotemporal overlap in spike times, for lag values between −500 −500 ms in increments of 1 ms. To control for variability in firing rate between recordings, the CCG values were normalized to the maximum value in this CCG vector.

## Code availability

Results were generated using code written in Matlab 2020b. All code is available at https://github.com/DanAkarca/MEA_generative_models

## Data availability

All data used in this study, along with documentation detailing each dataset, is openly available at https://zenodo.org/record/6109414#.Yid27y-l2J8

## ACKNOWLEDGEMENTS

This work was supported by the European Union through the European Research Council (ERC) Advanced Grant 694829 ‘neuroXscales’ and the corresponding proof-of-concept Grant 875609 ‘HD-Neu-Screen’, by the two Cantons of Basel through a Personalized Medicine project (PMB-01-18), granted by ETH Zurich, the Innosuisse Project 25933.2 PFLS-LS, the Swiss National Science Foundation under contract 205320_188910 / 1 and a Swiss Data Science Center project grant (C18-10). Danyal Akarca and Alexander Dunn are supported by the Medical Research Council Doctoral Training Programme. Danyal Akarca is supported by the Cambridge Trust Vice Chancellor’s Award Scholarship. Duncan Astle is supported by Medical Research Council Program Grant MC-A0606-5PQ41. Both Duncan Astle and Danyal Akarca are supported by The James S. McDonnell Foundation Opportunity Award. Congwei Wang and Marco Terrigno are supported by Roche postdoctoral fellowship program. Petra Vertes is a fellow of MQ:Transforming Mental Health (MQF17_24).

We thank Dr Martin Oeggerli for contributing the serial section electron microscopy image (**Figure 1a**), and the IT department at the MRC Cognition and Brain Sciences Unit, Cambridge, as well as the HPC team at ETH Zürich, for assistance with high performance computing.

## AUTHOR CONTRIBUTIONS

DA, AWED, DEA & MS conceived the project and wrote the manuscript. DA, AWED and MS contributed to all analyses provided in the manuscript. MS ran the processing of all neuronal data, including spike-sorting and functional connectivity inference. DA computed the generative network models, topological analyses of networks, topological fingerprints. AWED computed cellular firing and bursting analyses. MS, PJH, SR & MF recorded all neuronal data provided in the study. OP, SBM, SE, PEV provided computational and physiology overview, including STTC and gabazine expertise. AH & MS provided the engineering overview, particularly relating to HD-MEA recording. CW and MT cultured and derived human cerebral organoids.

