## Supplement for "Homophilic wiring principles underpin neuronal network topology *in vitro*"

### CONTENTS

#### *Supplementary Figures*


**Supplementary Figure 1.** Tracking neuronal networks at cellular resolution on high-density microelectrode arrays.

**Supplementary Figure 2.** Global topological measures of sparse rodent cultures over time.

**Supplementary Figure 3.** Distribution of inter-neuronal Euclidean distances across rodent and human datasets.

**Supplementary Figure 4.** Generative network modeling results for the sparse rodent primary cortical cultures.

**Supplementary Figure 5.** Alternative generative model procedures.

**Supplementary Figure 6.** Generative model fits recapitulate observed network statistics.

**Supplementary Figure 7.** Comparison of global network statistics across sparse and dense primary cortical rodent networks at DIV14.

**Supplementary Figure 8.** Generative network modeling results for the dense rodent primary cortical cultures.

**Supplementary Figure 9.** Homophilic generative mechanisms best account for local relationships in developing dense rodent neuronal cultures.

**Supplementary Figure 10.** Relationship between model energy and topological fingerprint dissimilarity.

**Supplementary Figure 11.** Chronic gabazine application to block $GABA_A$ receptor activity led to changes in spiking patterns.

**Supplementary Figure 12.** Chronic gabazine application to block $GABA_A$ receptor activity led to changes in functional network topology.

**Supplementary Figure 13.** Inferring inhibitory connections where there is negative spike transmission probability between two neuronal units according to their cross-correlation histogram of spiking counts.

**Supplementary Figure 14.** Generative model comparisons between control, gabazine-treated and washout sparse rodent PC cultures.

**Supplementary Figure 15.** Spiking dynamics in human monolayer and cerebral organoid cultures compared to rodent cultures.


#### *Supplementary Tables*


**Supplementary Table 1.** Overview of all the datasets used in the study.

**Supplementary Table 2.** A list of all the value $K_{i,j}$ terms that were included in the generative modeling, as given in the wiring equation.

**Supplementary Table 3.** Statistical comparisons of rodent 50k neuronal culture energy comparisons across generative rules.

**Supplementary Table 4.** Statistical comparisons of rodent 100k neuronal culture energy comparisons across generative rules.

**Supplementary Table 5.** Statistical comparisons of rodent 50k neuronal culture topological fingerprint dissimilarity comparisons across generative rules.

**Supplementary Table 6.** Statistical comparisons of rodent 100k neuronal culture topological fingerprint dissimilarity comparisons across generative rules.

**Supplementary Table 7.** Statistical comparisons of human iPSC neuronal culture (DIV28 glutamatergic neurons, DIV28 motor neurons and DIV28 dopaminergic neurons) and human cerebral organoids energy comparisons across generative rules.

**Supplementary Table 8** Overview of used antibodies.

### SUPPLEMENTARY FIGURES

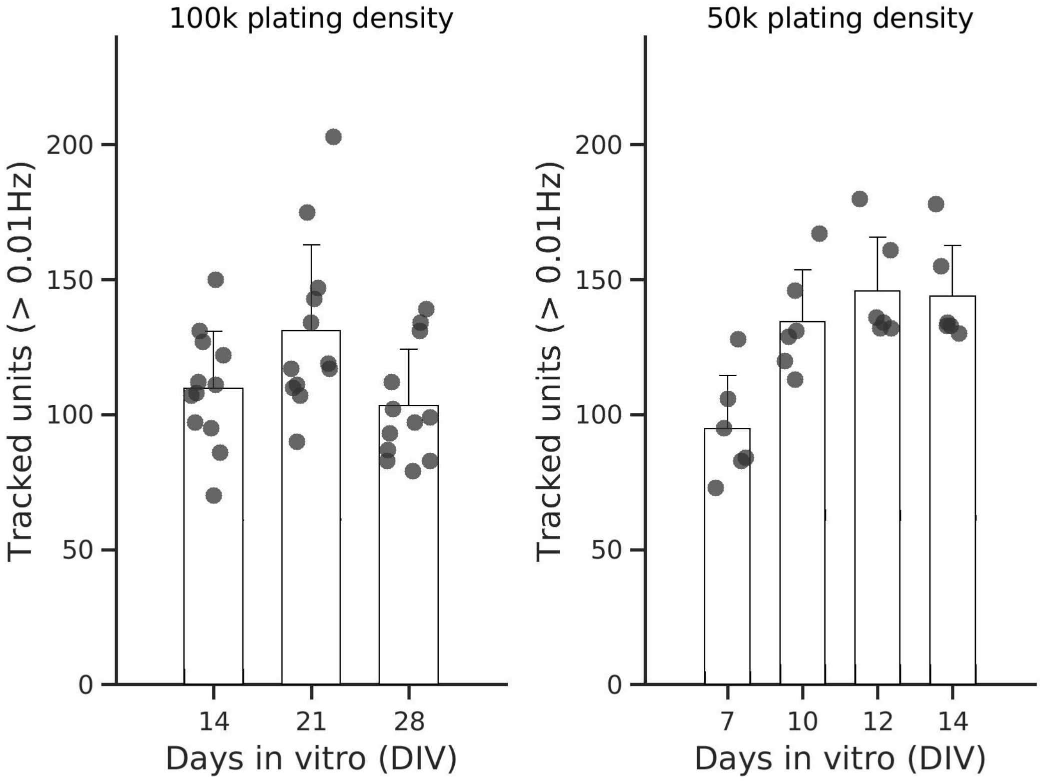


***Supplementary Figure 1.***

**Tracking neuronal networks at cellular resolution on high-density microelectrode arrays.**

Bar plots depicting the tracking result for the rodent data with 100k (left) and 50k (right) plating density per high-density microelectrode array (HD-MEA).

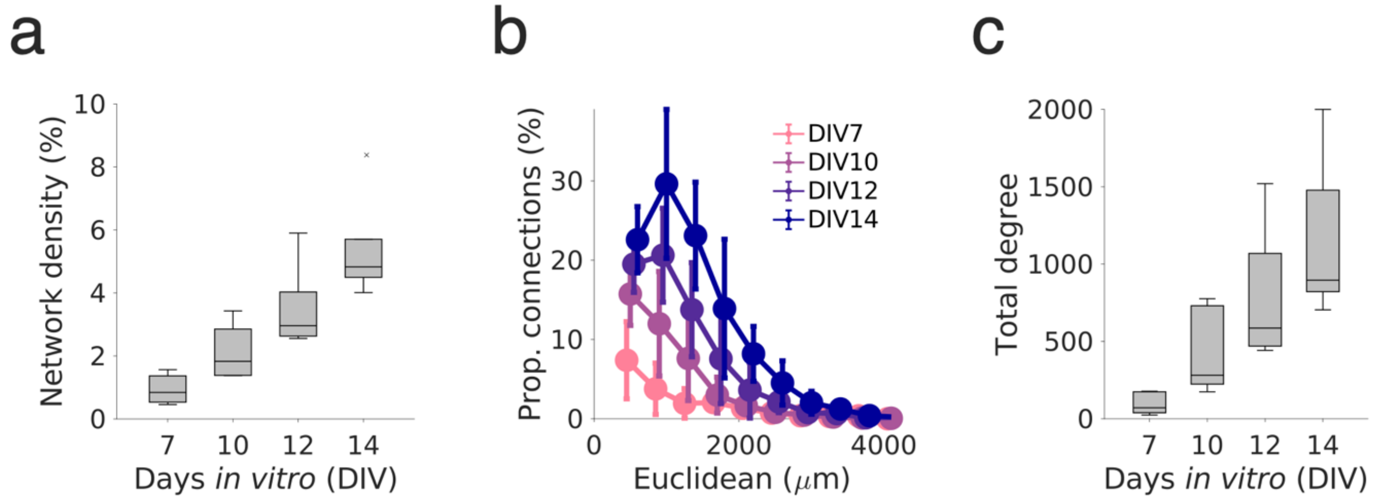


*Supplementary Figure 2.*

**Global topological measures of sparse rodent cultures over time**.

**a** Panel depicts the statistically inferred network density for the sparse (50,000 cell per well) primary cortical (PC) neuronal networks until days in vitro (DIV)14. **b** Proportion of extant connection by distance and grouped by DIV7 (pink), DIV10 (light purple), DIV12 (dark purple) and DIV14 (dark blue). **c** The total degree for sparse PC networks across development.

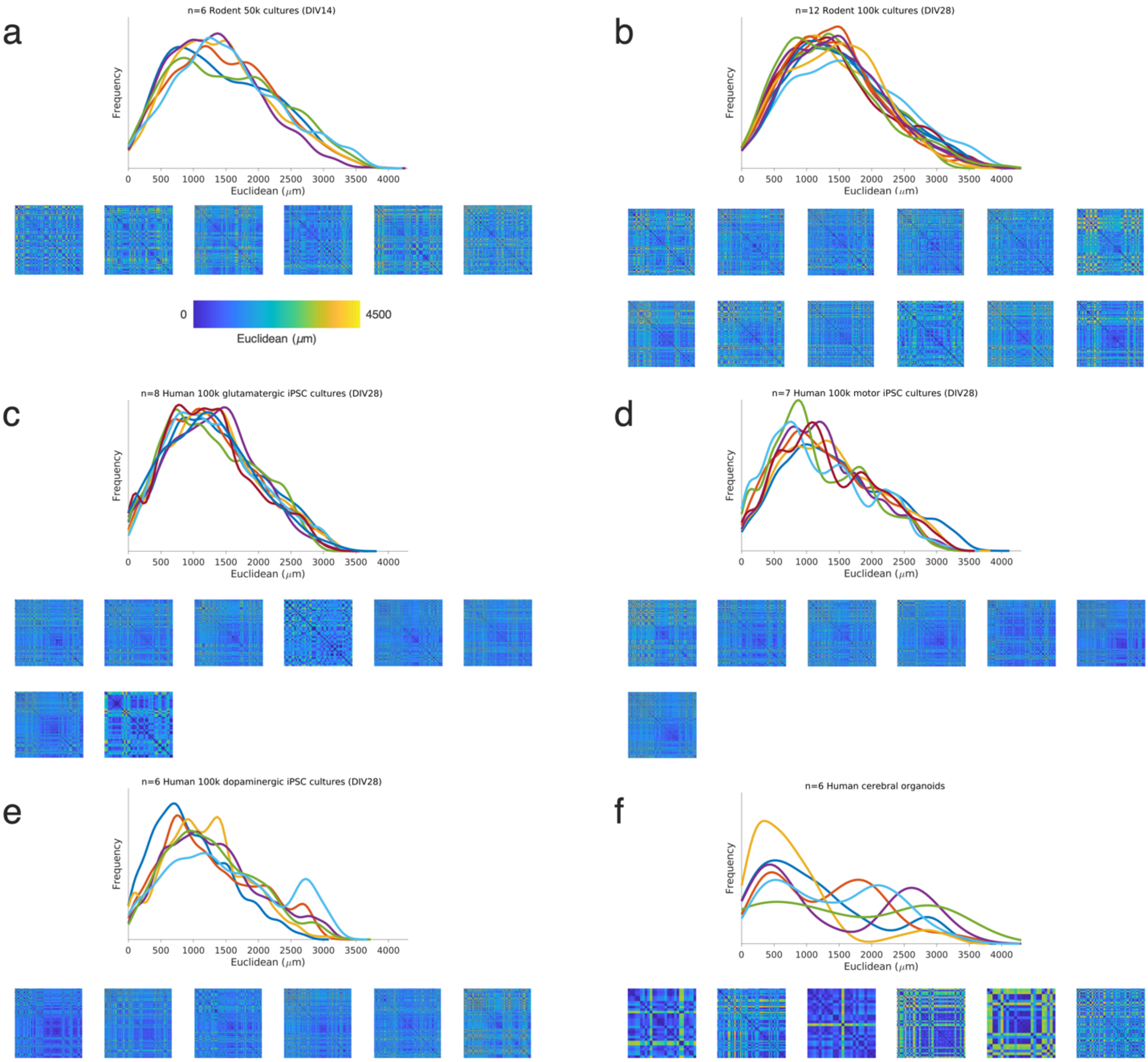


***Supplementary Figure 3.***

**Distribution of inter-neuronal Euclidean distances across rodent and human datasets.**

**a** Density plot of inter-neuronal Euclidean distances for each culture. Note, each matrix was used as the $D_{i,j}$ term for the generative model constructed for each time point of that culture. Panel **a** depicts Euclidean distances for the sparse PC rodent networks (50,000 neurons per well); the left panel shows the overall distributions; the right panel shows the individual distance matrices. **b** Euclidean distances for the dense PC rodent networks (100,000 neurons per well). Panels **c-e** show Euclidean distance distribution across the iPSC-derived human neuron lines at DIV28 (**c**, glutamatergic; **d**, motor and **e**, dopaminergic neurons). **f** Euclidean distances for the human cerebral organoid recordings.

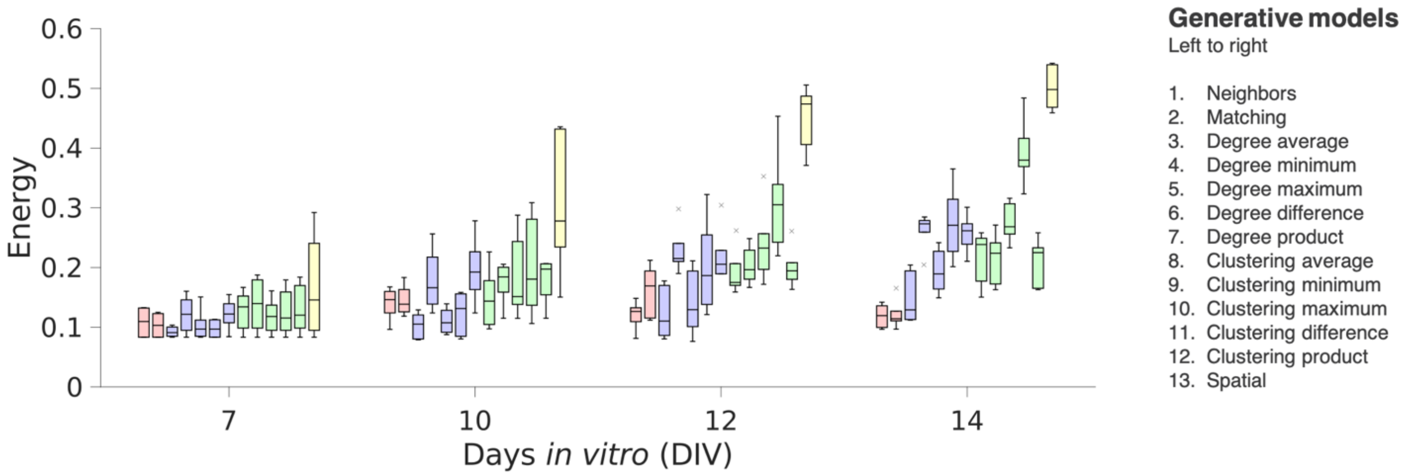


***Supplementary Figure 4.***

**Generative network modeling results for the sparse rodent primary cortical cultures.**

Generative model fits (energy), including all 13 wiring models, for the sparse primary rodent networks (50,000 cells per well; n=6 cultures) across development. Each boxplot presents the median and IQR. Outliers are demarcated as small black crosses, and are those which exceed 1.5x the interquartile range away from the top or bottom of the box. Generative model performance over time according to the energy equation. In each box, the energy of the top n=1 performing simulations are shown.

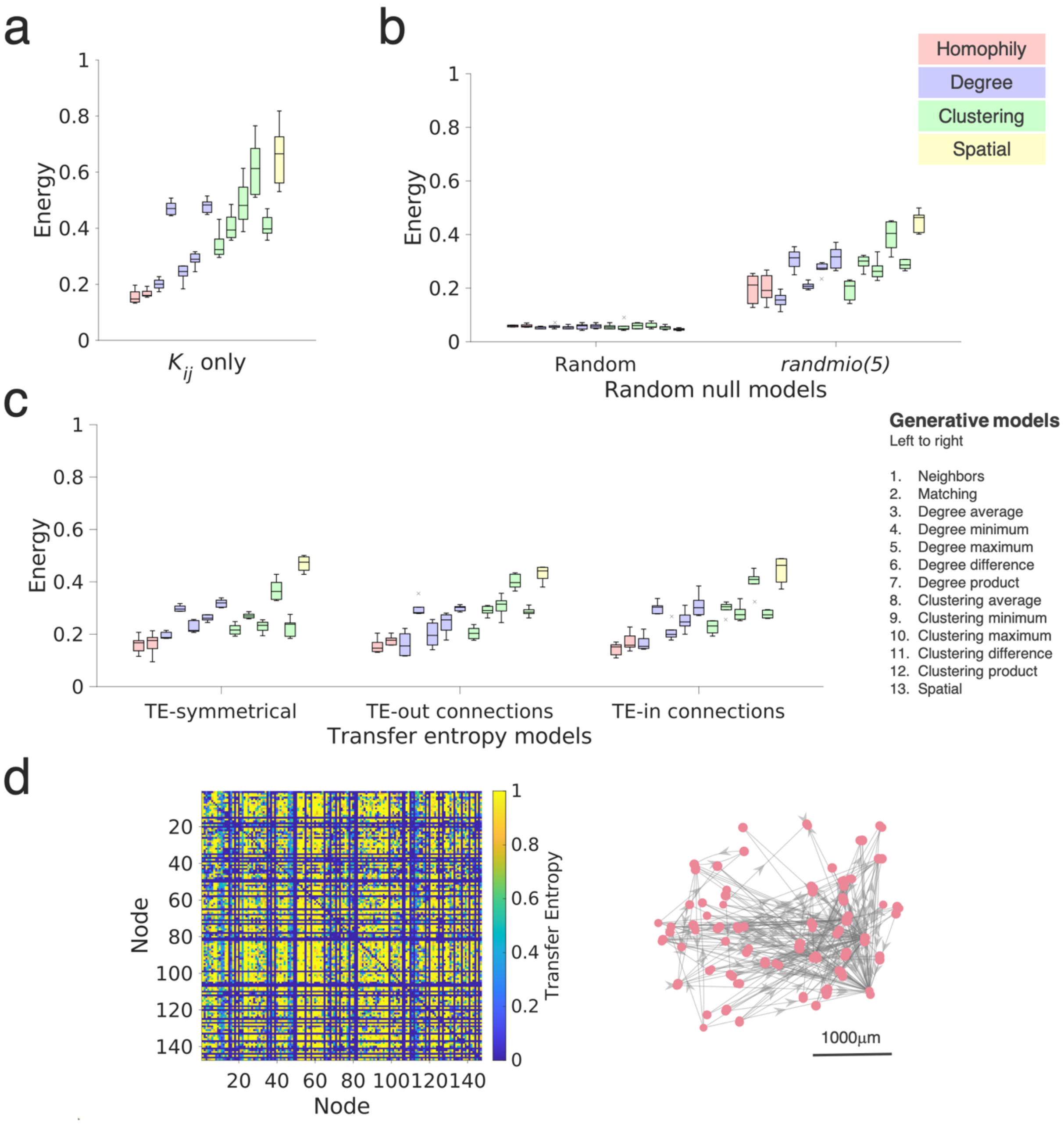


***Supplementary Figure 5.***

**Alternative generative model procedures.**

Generative model fits (energy), including all 13 wiring models, for 50k-plated primary rodent networks (50,000 cells per well; n=6 cultures) at DIV14. **a** Generative models fit only using the wiring term ($K_{i,j}$) when forming networks. **b** Random null models. Completely randomized networks (size- and density-matched, left) and networks which have been partially randomized via the *randmio_und()* function, using the Maslov-Sneppen algorithm, which rewires each edge approximately five times. **c** Inferred transfer entropy (TE) networks. As the TE is a directed graph, we show the results for the symmetrized network (left); out-connections (middle) and in-connections (right). For each graph, the boxplot presents the median and IQR. Outliers are demarcated as small black crosses, and are those which exceed 1.5x the interquartile range away from the top or bottom of the box. Generative model performance over time according to the energy equation. In each box, the energy of top n=1 performing simulation is shown. **d** A visualization of the transfer entropy matrix (left) of a single empirical network (right).

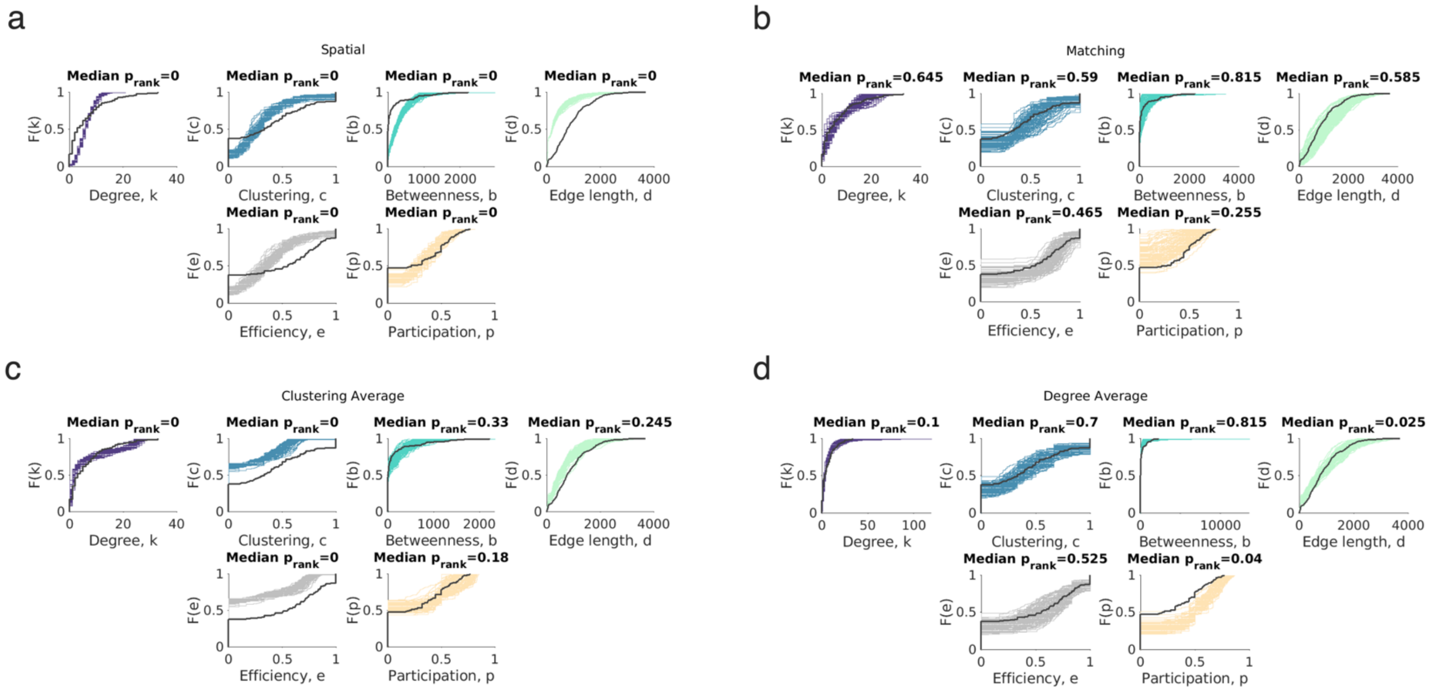


***Supplementary Figure 6.***

**Generative model fits recapitulate observed network statistics.**

**a** Cumulative density functions (CDFs) of the top=99 simulations (of the total 20,000 simulations) for the best performing generative models in each class (**a**, spatial. **b**, matching. **c**, clustering average. **d**, degree average) across the four statistics included in the energy equation (top four panels) and two additional metrics - the local efficiency and participation coefficient - not included in the energy equation (bottom two panels). For visualization, we show only the solutions for a single sparse rodent PC culture. p values were computed using the Monte-Carlo bootstrapping procedure outlined in **Methods; *Generative network models***.

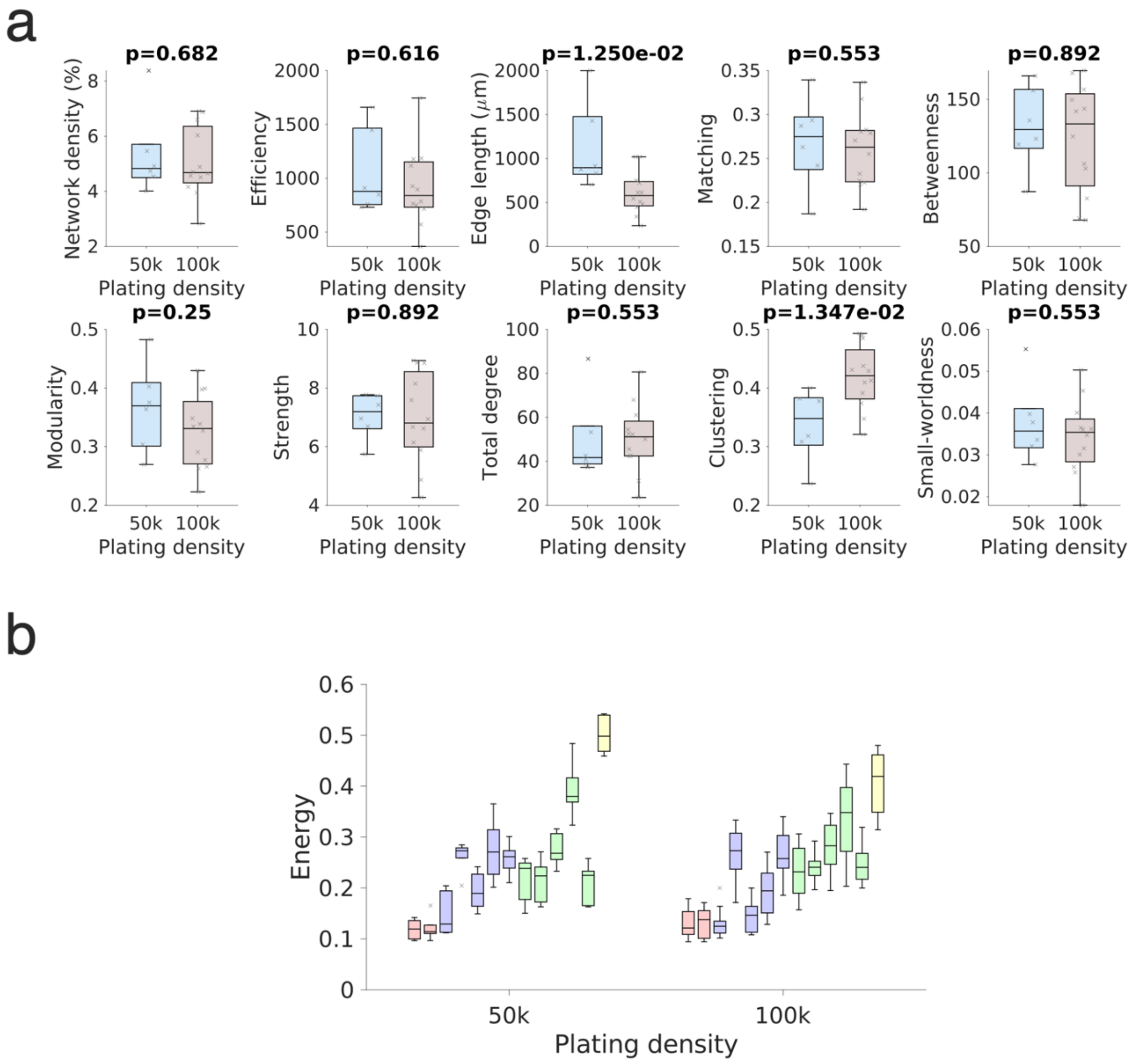


*Supplementary Figure 7.*

**Comparison of global network statistics across sparse and dense primary cortical rodent networks at DIV14.**

**a** Comparisons were computed for network density, efficiency, edge length, matching, betweenness, modularity, strength, total degree, clustering and small worldness. For each comparison we provide the p-value computed from a Mann-Whitney U test. Sparse networks were plated at 50,000 cells per well, dense networks were plated at 100,000 cells per well. **b** Direct comparisons of the generative model fits (energy), including all 13 wiring models, for the 50k-plated (left) versus 100k-plated (right) primary rodent networks at DIV14. The boxplot presents the median and IQR. Outliers are demarcated as small black crosses, and are those which exceed 1.5x the interquartile range away from the top or bottom of the box. Generative model performance over time according to the energy equation. In each box, the energy of top n=1 performing simulation is shown.

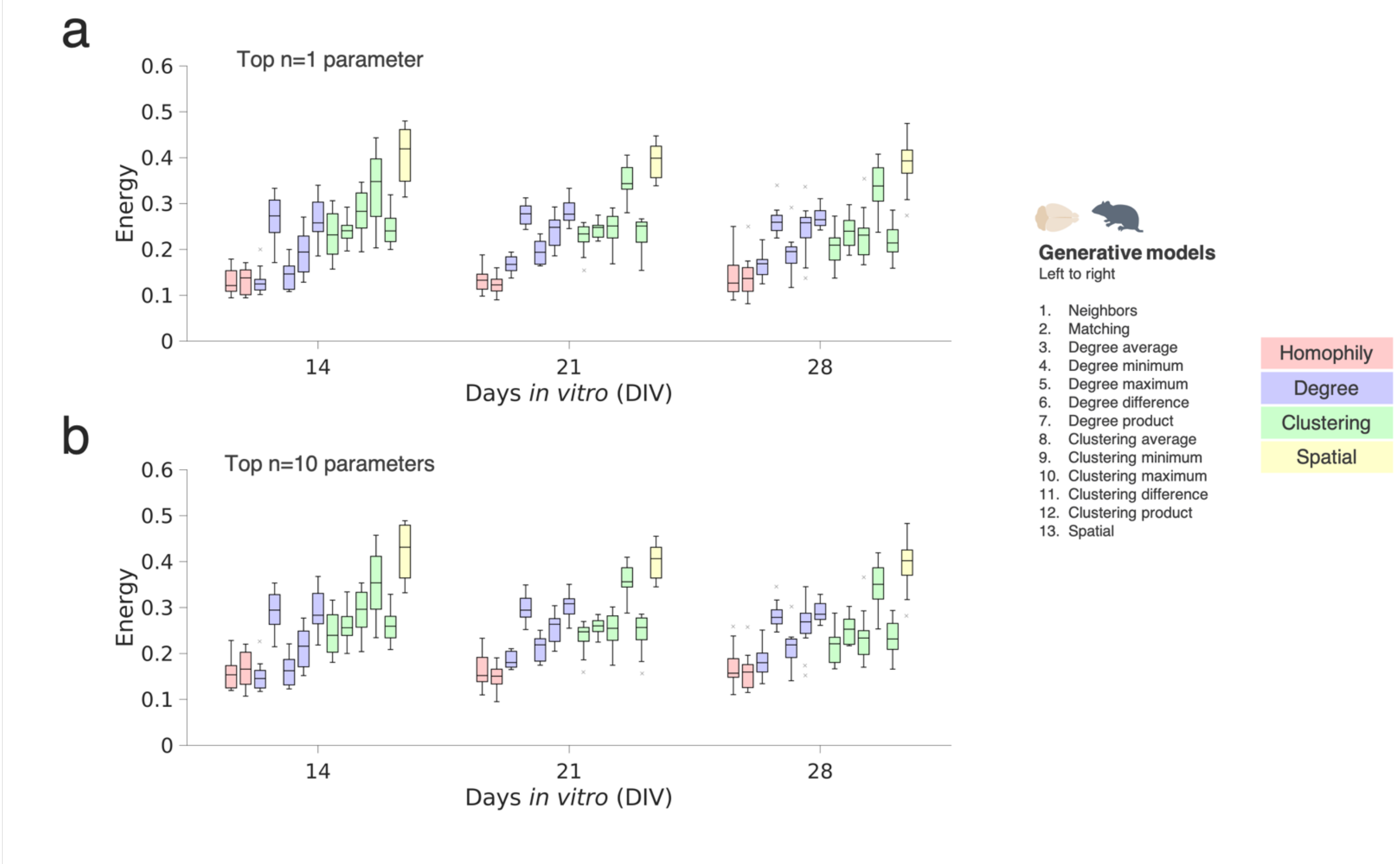


***Supplementary Figure 8.***

**Generative network modeling results for the dense rodent primary cortical cultures.**

Generative model fits (energy), including all 13 wiring models, for the dense primary rodent networks (100,000 cells per well; n=12 cultures) across development. **a** Top performing n=1 parameter combination. **b** Top performing n=10 parameter combinations. Each boxplot presents the median and IQR. Outliers are demarcated as small black crosses, and are those which exceed 1.5x the interquartile range away from the top or bottom of the box. Generative model performance over time according to the energy equation. In each box, the energy of the top n=1 performing simulations are shown.

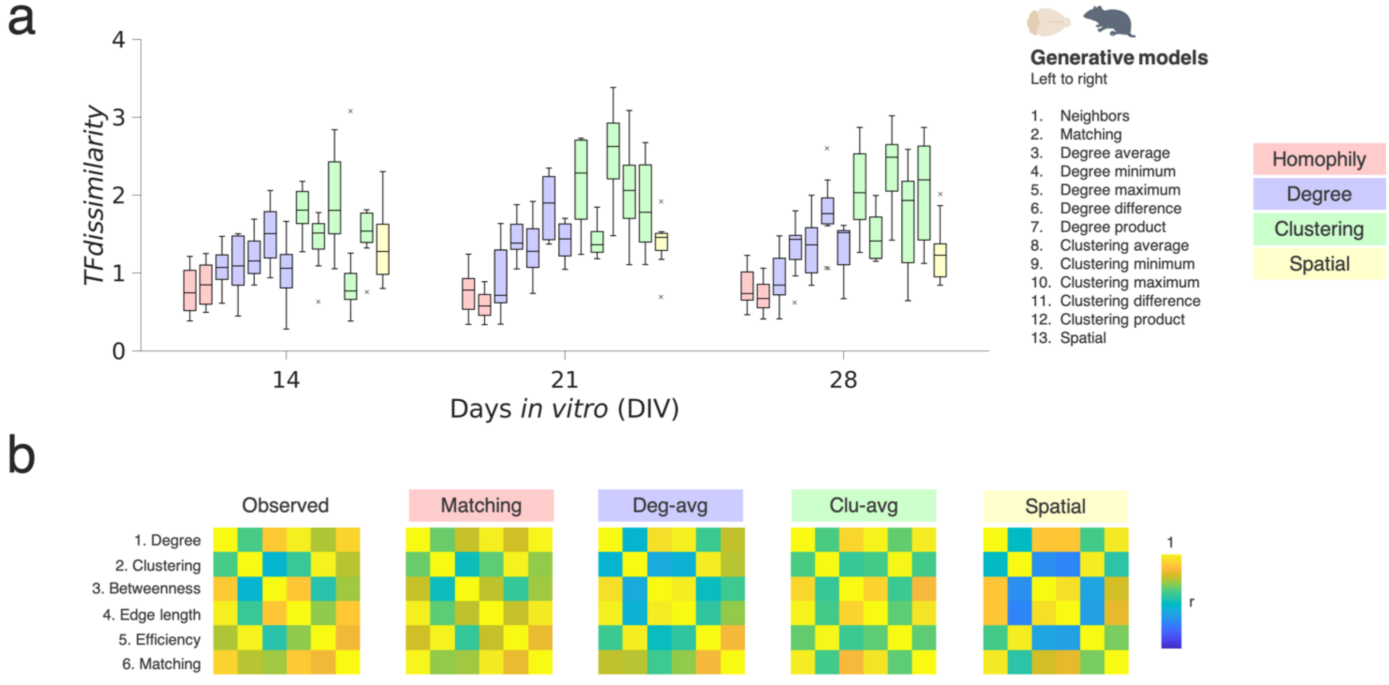


*Supplementary Figure 9.*

**Homophilic generative mechanisms best account for local relationships in developing dense rodent neuronal cultures.**

**a** Homophily generative models produce the lowest *TFdissimilarity* across all time-points (DIV14, 21 and 28), suggesting that it can reconstruct local connectivity patterns of in *vitro* neuronal networks. In total, there are n=12 data points (one per culture) shown in each of the 13 boxplots. Boxplots presents the median and IQR. Outliers are demarcated as small gray crosses, and are those which exceed 1.5 times the interquartile range away from the top or bottom of the box. **b** A visualization of the averaged topological organization matrix for the observed (left), matching (middle left), degree-average (middle), clustering-average (middle right), and spatial (right) models at DIV28.

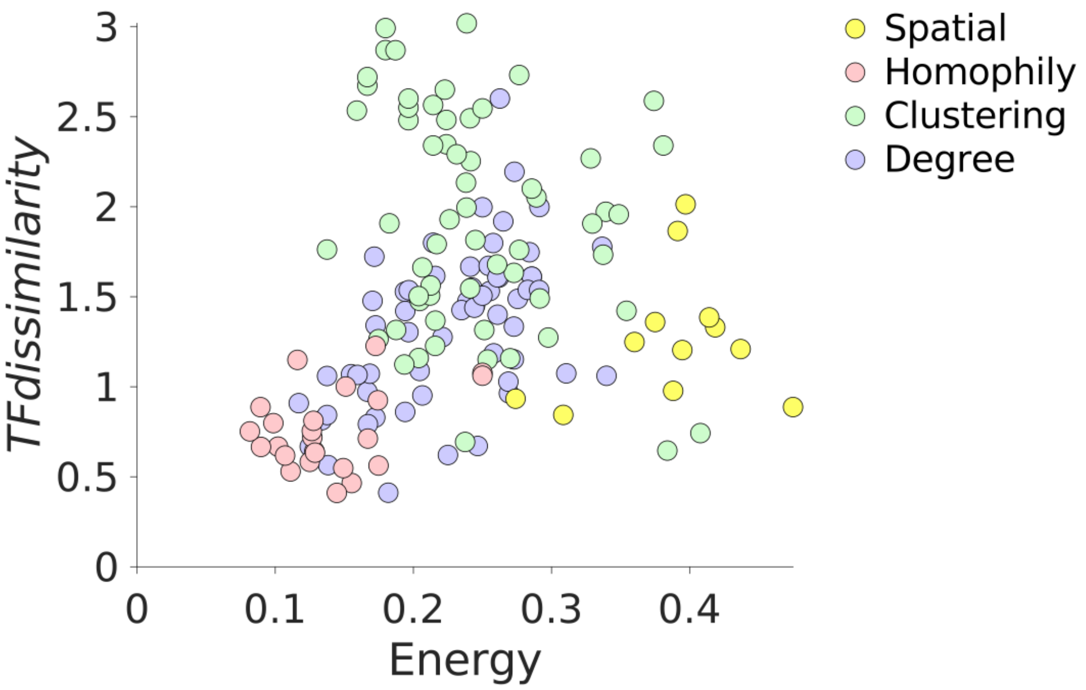


***Supplementary Figure 10.***

**Relationship between model energy and topological fingerprint dissimilarity in developing dense rodent neuronal cultures.**

Using dense PC rodent network (100,000 cells per MEA), we show n=78 data points (n=6 cultures x n=13 generative model simulations) corresponding to the top n=1 performing simulation's energy and its topological fingerprint dissimilarity performances at DIV14. Distributions are plotted for each, showing homophily to achieve the best fits in both (bottom left).

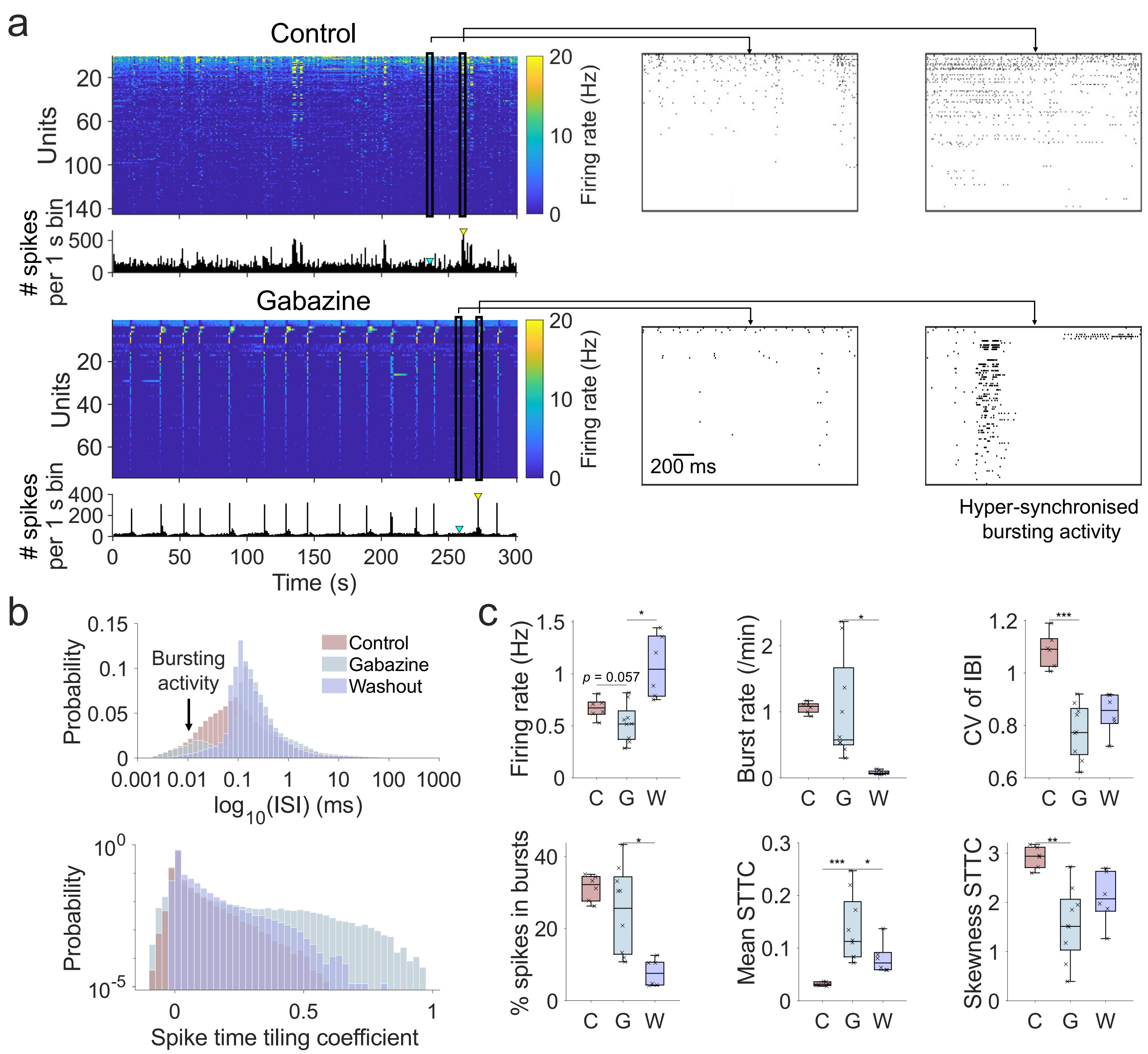


***Supplementary Figure 11.***

**Chronic gabazine application to block $GABA_A$ receptor activity led to changes in spiking patterns.**

**a** Representative spike train raster plots from a control (left) and a gabazine-treated culture (right); the panel below shows the representative population activity vectors (activity is aggregated 1 s bins). Panels on the bottom: example close-ups of 1 s time windows taken from time bins comprising either median network activity (left, indicated by cyan triangle) or peak network activity (right, indicated by yellow triangle). **b** A logarithmic histogram of interspike intervals (ISIs; upper panel) showing that gabazine relatively increases the proportion of very short ISIs and long ISIs, giving a tri-modal distribution, equivalent to periods of relative quiescence followed by periods of fast bursting. The spike time tiling coefficient (STTC) distribution (probability plotted on a log10 scale for visualization) shows gabazine-treated cultures had a far greater number of strongly correlated units compared to controls in which there were far fewer strong functional connections. **c** Cultures treated with Gabazine showed decreased mean firing rate (Hz) of cellular activity compared to controls, but increased and more regular spiking and bursting activity (reduced coefficient of variation in inter-spike and inter-burst intervals). Gabazine treatment also led to much stronger edge weights (STTC values) and less positively skewed edge weight distributions (less positive skewness of STTC distribution)—this skewness in controls reflects the presence of high strength (hub) nodes. Asterisks, *, ** and *** indicated $p < 0.05$, 0.01 and 0.001, respectively.

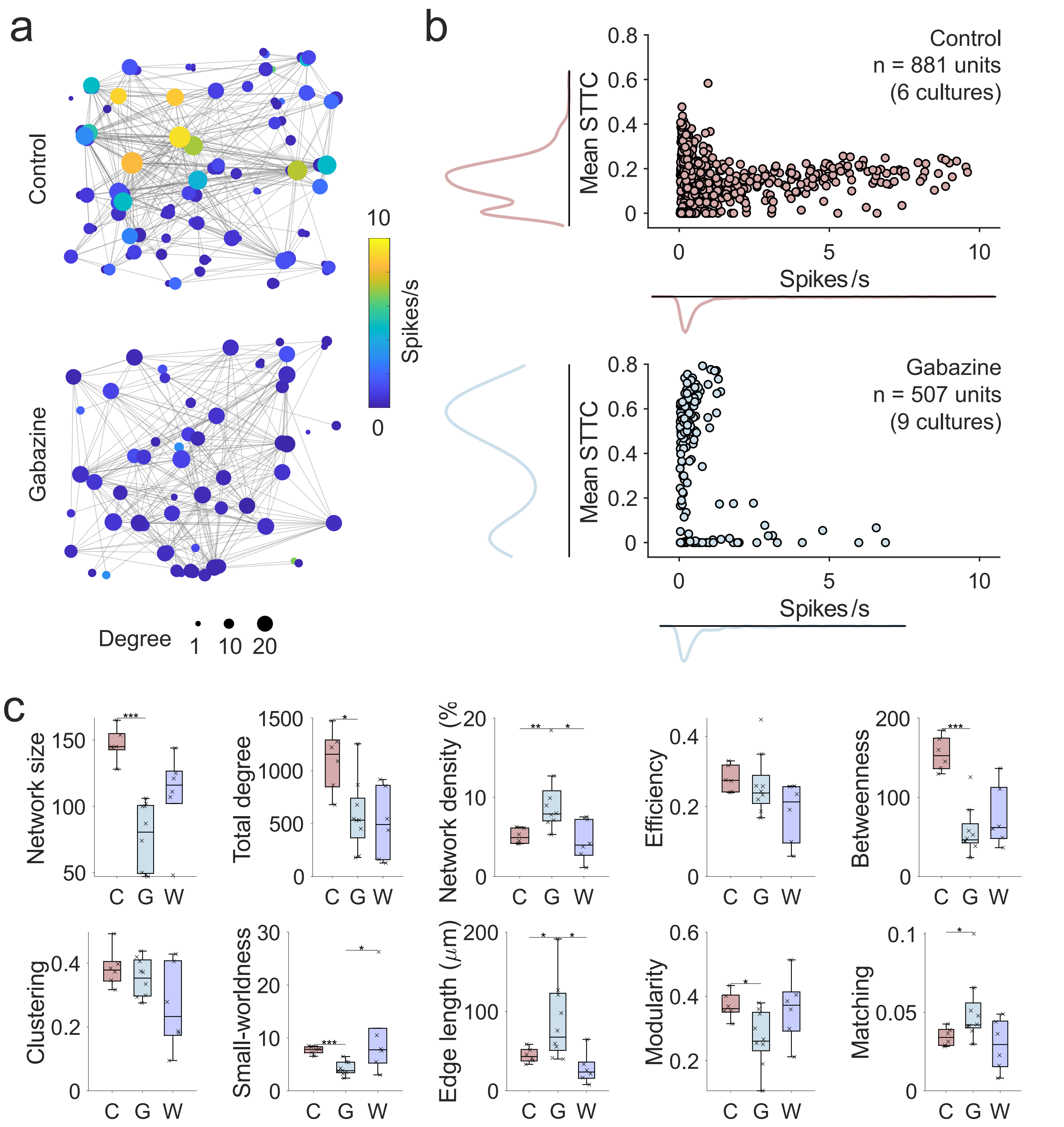


*Supplementary Figure 12.*

**Chronic gabazine application to block $GABA_A$ receptor activity led to changes in functional network topology.**

**a** Representative examples of topological network plots from a 14-days-in-vitro rodent cortical culture (DIV; upper panel) and a culture treated with gabazine from DIV1 (lower below). Each node (colored circles) represents the spatial location of the soma of a putative neuronal unit detected through spike sorting. Each edge (grey lines) represents a significant correlation in spiking activity between a pair of nodes. Functional connections were inferred using the spike timing tiling coefficient (STTC). The size of each node is proportional to the number of significant functional connections; it's color reflects its spike rate. **b** Each dot represents a single neuronal unit. For both control cultures (upper; brown) and gabazine-treated cultures (lower; blue) there was no clear linear relationship between spike rate and mean STTC (average weight of all significant edges of a given node). Each group had similar firing rate distributions, though very different average functional connection weight distributions with gabazine-treated units being hyperconnected compared to controls. **c** A range of global topological metrics inferred from functional connectivity graphs of control (C) and Gabazine (G) treated PC cultures. For each comparison we provide the p-value computed from a Mann-Whitney U test. Asterisks, *, ** and *** indicated $p < 0.05$, 0.01 and 0.001, respectively.

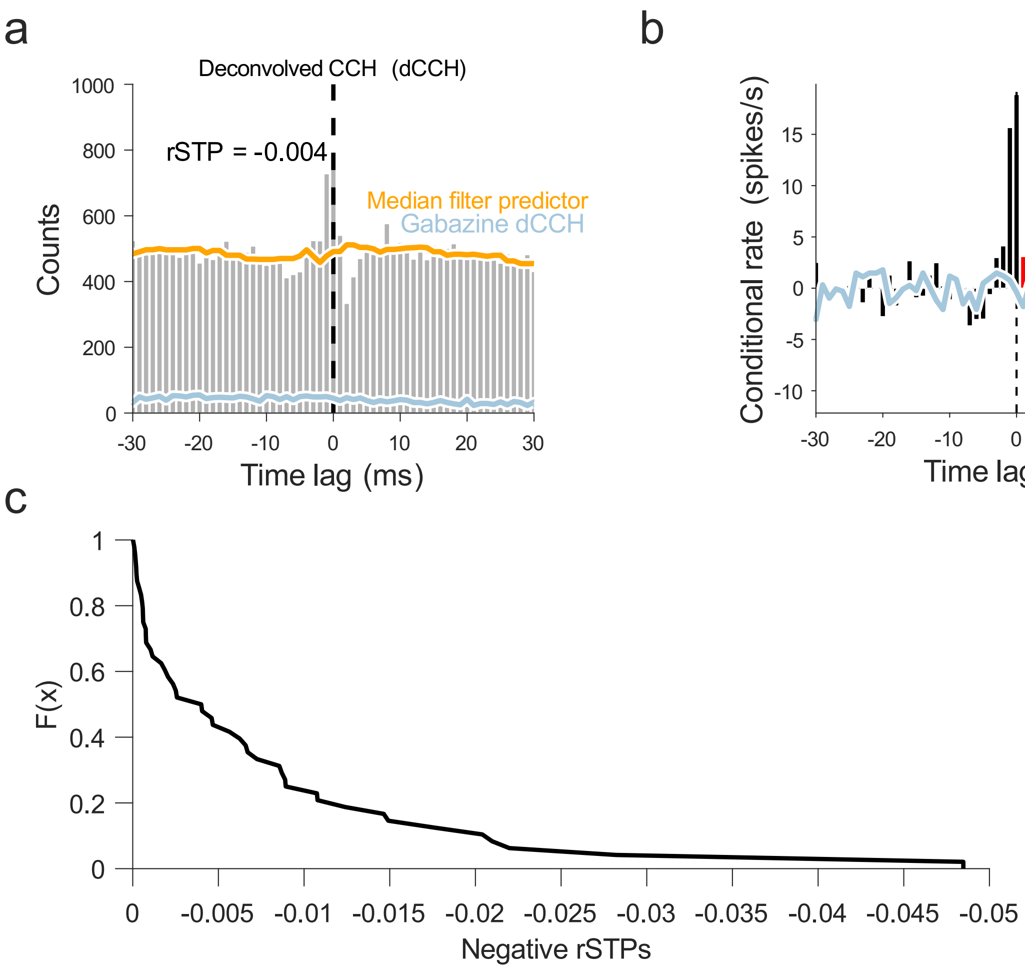


***Supplementary Figure 13.***

**Inferring inhibitory connections where there is negative spike transmission probability between two neuronal units according to their cross-correlation histogram of spiking counts.**

**a** An example deconvolved cross-correlation histogram of overlapping spiking activity across time lags tested between two neurons in a culture treated with gabazine then washed out. Median spike count prediction across time lag values is indicated by the orange line. **b** The same two neuronal units with spike count relative to the prediction plotted. Red bars indicate where the spike count was less than predicted in neuron *j* between 0 – 5 ms after neuron *i* fired, thus indicating a putative inhibitory connection. **c** Cumulative distribution of negative ratio of spike transmission probability across the culture showing the majority of connections are not inhibitory (rSTP ≈ 0), while around 20 % show some inhibition (rSTP < -0.01).

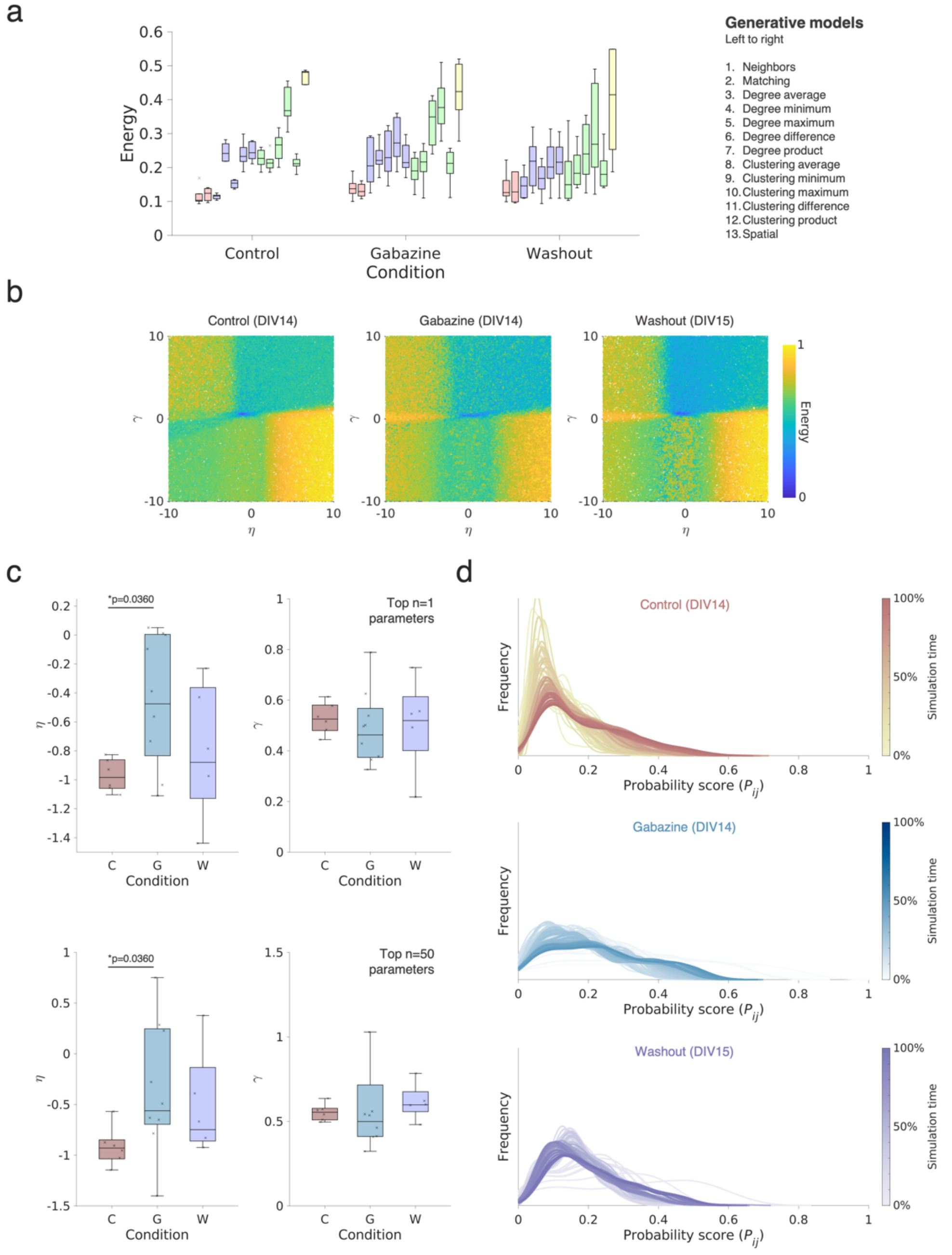


*Supplementary Figure 14.*

**Generative model comparisons between control, gabazine-treated and washout sparse rodent PC cultures.**

**a** Direct comparisons of the generative model fits (energy), including all 13 wiring models, for control (left) versus gabazine (middle) and washout (right) cultures. The boxplot presents the median and IQR. Outliers are demarcated as small black crosses, and are those which exceed 1.5x the interquartile range away from the top or bottom of the box. Generative model performance over time according to the energy equation. In each box, the energy of top n=1 performing simulation is shown. **b** The energy landscapes of the matching generative rule for control (left) versus gabazine (middle) and washout (right) cultures. **c** Wiring parameters of control (left; colored in red), gabazine (middle; colored in blue) and washout (right; colored in purple) derived from the top n=1 (top) and n=50 (bottom) performing matching simulation in terms of the wiring equation are shown. **d** The probability distributions for control (top), gabazine (middle) and washout (bottom) networks were computed and scaled at 1% increments throughout the developmental course of the simulations. These distributions were averaged over time to form the mean probability distributions shown in **Figure 6d**.

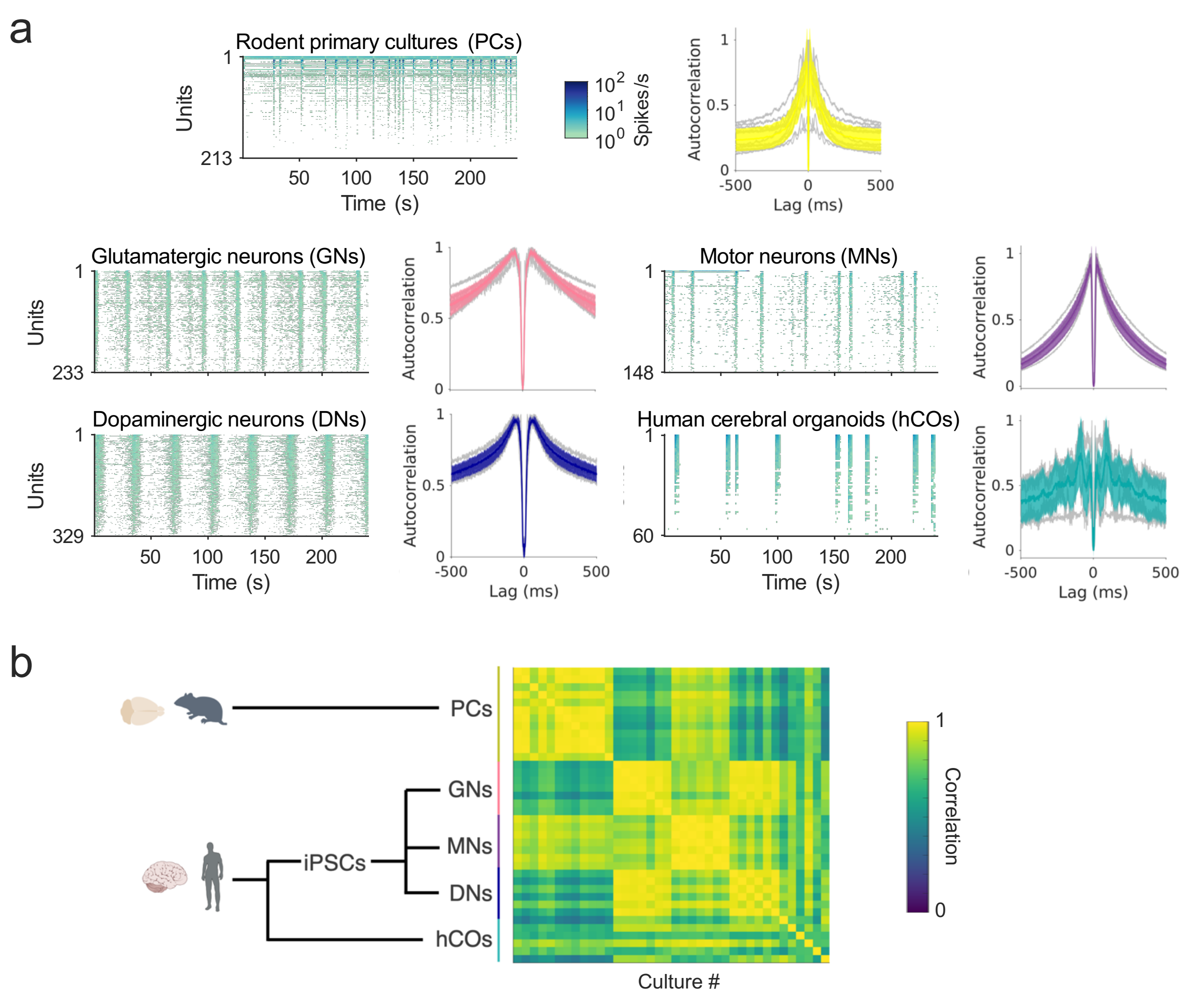


***Supplementary Figure 15.***

**Spiking dynamics in human monolayer and cerebral organoid cultures compared to rodent cultures.**

**a** Temporal raster plots indicate the firing rates (spikes/s) of each cell line over consecutive 1 s time bins through 5 minutes of recording. Motor neuron patterns more closely resemble rodent cortical networks. Both glutamatergic and dopaminergic iPSC cultures showed highly regular network bursting activity, though at differing frequencies. Human cerebral organoid networks had fewer units but had periods of much higher firing rates. For each culture type, autocorrelograms of spiking activity are plotted to the right of each temporal raster plot. Autocorrelograms were constructed by first cumulatively concatenating spike times across all units in a recording into a single vector of spike times. The count of synchronous spikes across the network (in 1 ms bins) was quantified across a range of time lags (0 lag was excluded). This indicates the degree of coactivation at various latencies. In all cell culture types, there was much higher spiking synchrony at shorter latencies indicated by peaks close to 0 ms lag. However, rodent primary cultures (PCs) and human motor neurons (MNs) showed sharp declines in autocorrelation at longer latencies indicating a strong preference for short latency, high frequency coactivation. Human glutamatergic (GNs) and dopaminergic (DNs) cultures had much greater autocorrelation at lag values further from 0. This indicates coactivation at longer latencies and oscillatory network activity at slower frequencies. Human cerebral organoids (hCOs) were much more variable. **b** The autocorrelation vectors (autocorrelation at each lag value) across all cultures were correlated and a high level of concordance within culture sources in terms of latency preference can be seen, with moderate correlations between rodent PCs and human MNs.

#### SUPPLEMENTARY TABLES

| Cell line | No. of units | Recording time point | Plating density | Samples (n) |
|---|---|---|---|---|
| Primary rodent cortex (sparse) | 129.7 ± 27.9 | DIV 7, 10, 12, 14 | 50,000 | 6 |
| Primary rodent cortex (dense) | 114.7 ± 27.4 | DIV 14/15, 21, 28 | 100,000 | 12 |
| Human motor neurons | 210.7 ± 24.7 | DIV 28 | 100,000 | 7 |
| Human glutamatergic neurons | 191.6 ± 69.1 | DIV 28 | 100,000 | 8 |
| Human dopaminergic neurons | 211.2 ± 55.9 | DIV 28 | 100,000 | 6 |
| Human cerebral organoids | 47.3 ± 27.8 | 120 days | n/a | 6 (slices),<br>3 organoids |

***Supplementary Table 1.***

**Overview of all the datasets used in the study.**

Number of unit results are expressed as the mean ± SD of the respective cell line.

| Name | $K_{i,j}$ |
|---|---|
| Neighbors | $\sum_l A_{il}A_{jl}$ |
| Matching | $\frac{\lvert N_{i/j} \cap N_{j/i} \rvert}{\lvert N_{i/j} \cup N_{j/i} \rvert}$ |
| Clustering Average | $\frac{c_i}{2} + \frac{c}{2}$ |
| Clustering Difference | $\lvert c_i - c_j \rvert$ |
| Clustering Maximum | $\max(c_i, c_j)$ |
| Clustering Minimum | $\max(c_i, c_j)$ |
| Clustering Product | $c_i c_j$ |
| Degree Average | $\frac{k_i}{2} + \frac{k_j}{2}$ |
| Degree Difference | $\lvert k_i - k_j \rvert$ |
| Degree Maximum | $\max(k_i, k_j)$ |
| Degree Minimum | $\max(k_i, k_j)$ |
| Degree Product | $k_i k_j$ |
| Spatial | 1 |

***Supplementary Table 2.***

**List of all value $K_{i,j}$ terms that were included in the generative modeling, as given in the wiring equation.**

*A* is the binary adjacency matrix, c is the local clustering coefficient, *k* is the node degree and $N_{i/j}$ represents the neighbors of node *i*, excluding node *j*. Note that the spatial model enforces $K_{i,j}$ =1, which means that the value term has no effect on the generative process.

| Rule A | Rule B | DIV 7 | | DIV10 | | DIV12 | | DIV14 | |
|---|---|---|---|---|---|---|---|---|---|
| | | ANOVA p=0.00134 | Cohen's *d* | DIV10 p=1.51e-07 | Cohen's *d* | ANOVA p=3.43e-15 | Cohen's *d* | ANOVA p=3.26e-17 | Cohen's *d* |
| Homophily | Degree | 1 | 0.0482 | 1 | -0.00443 | 0.222 | 0.583 | 4.11e-05 | 1.46 |
| Homophily | Clustering | 0.286 | 0.703 | 0.29 | 0.704 | 0.000528 | 1.52 | 7.01e-08 | 2.94 |
| Homophily | Spatial | 0.00592 | 0.926 | 1.47e-06 | 1.38 | 3.77e-09 | 4.83 | 3.77e-09 | 12 |
| Degree | Clustering | 0.127 | 0.6 | 0.0909 | 0.585 | 0.0242 | 0.646 | 0.16 | 0.42 |
| Degree | Spatial | 0.00219 | 0.675 | 9.12e-08 | 1.5 | 3.77e-09 | 6.3 | 3.77e-09 | 15.3 |
| Clustering | Spatial | 0.0802 | 0.496 | 2.25e-05 | 1.71 | 3.81e-09 | 4.44 | 3.80e-09 | 3.99 |

***Supplementary Table 3.***

**Statistical comparisons of rodent 50k neuronal culture energy comparisons across generative rules.**

For each test, we quote the ANOVA p-value across the generative rules and the corresponding Cohen's *d* if the ANOVA was significant at $p<0.05$. A positive Cohen's *d* reflects that Rule A has a smaller energy than Rule B, reflecting a better fit. Generative rules have been binned across the generative model class.

| Rule A | Rule B | DIV14 | | DIV21 | | DIV28 | |
|---|---|---|---|---|---|---|---|
| | | ANOVA p=7.55e-27 | Cohen's *d* | ANOVA p=7.04e-32 | Cohen's *d* | ANOVA p=1.27e-23 | Cohen's *d* |
| Homophily | Degree | 1.53e-05 | 0.843 | 3.77e-09 | 1.52 | 4.42e-09 | 1.19 |
| Homophily | Clustering | 3.77e-09 | 2.37 | 3.77e-09 | 3.05 | 3.77e-09 | 1.44 |
| Homophily | Spatial | 3.77e-09 | 3.33 | 3.77e-09 | 8 | 3.77e-09 | 3.1 |
| Degree | Clustering | 3.43e-08 | 0.745 | 0.0146 | 0.397 | 0.257 | 0.281 |
| Degree | Spatial | 3.77e-09 | 2.8 | 3.77e-09 | 4.14 | 3.77e-09 | 3.59 |
| Clustering | Spatial | 3.78e-09 | 3.92 | 3.77e-09 | 2.32 | 3.77e-09 | 2.4 |

***Supplementary Table 4.***

**Statistical comparisons of rodent 100k neuronal culture energy comparisons across generative rules.**

For each test, we quote the ANOVA p-value across the generative rules and the corresponding Cohen's *d* if the ANOVA was significant at $p<0.05$. A positive Cohen's *d* reflects that Rule A has a smaller energy than Rule B, reflecting a better fit. Generative rules have been binned across the generative model class.

| Rule A | Rule B | DIV7 | | DIV10 | | DIV12 | | DIV14 | |
|---|---|---|---|---|---|---|---|---|---|
| | | ANOVA p=0.0847 | Cohen's *d* | ANOVA p=2.6e-09 | Cohen's *d* | ANOVA p=1.81e-05 | Cohen's *d* | ANOVA p=3.9e-07 | Cohen's *d* |
| Homophily | Degree | n/a | n/a | 0.725 | -0.272 | 0.0762 | 1.15 | 0.00191 | 1.34 |
| Homophily | Clustering | n/a | n/a | 0.000167 | 2.39 | 6.35e-05 | 3.58 | 1.37e-07 | 2.59 |
| Homophily | Spatial | n/a | n/a | 1 | 0.00263 | 0.000996 | 1.39 | 0.0952 | 0.818 |
| Degree | Clustering | n/a | n/a | 5.55e-09 | 1.55 | 0.0189 | 0.544 | 0.00749 | 0.726 |
| Degree | Spatial | n/a | n/a | 0.851 | 0.704 | 0.0615 | 0.733 | 0.995 | -0.0648 |
| Clustering | Spatial | n/a | n/a | 0.006 | -0.967 | 0.849 | 0.291 | 0.147 | -0.515 |

***Supplementary Table 5.***

**Statistical comparisons of rodent 50k neuronal culture topological fingerprint dissimilarity comparisons across generative rules.**

For each test, we quote the ANOVA p-value across the generative rules and the corresponding Cohen's *d* if the ANOVA was significant at $p<0.05$. A positive Cohen's *d* reflects that Rule A has a smaller energy than Rule B, reflecting a better fit. Generative rules have been binned across the generative model class.

| Rule A | Rule B | DIV14 | | DIV21 | | DIV28 | |
|---|---|---|---|---|---|---|---|
| | | ANOVA p=1.01e-08 | Cohen's *d* | ANOVA p=1.6e-21 | Cohen's *d* | ANOVA p=2.09e-18 | Cohen's *d* |
| Homophily | Degree | 0.00492 | 0.996 | 7.87e-09 | 1.27 | 3.89e-06 | 1.19 |
| Homophily | Clustering | 4.37e-09 | 1.79 | 3.77e-09 | 2.87 | 3.77e-09 | 2.76 |
| Homophily | Spatial | 0.00332 | 0.869 | 0.000125 | 1.58 | 0.0138 | 1.02 |
| Degree | Clustering | 0.000195 | 0.519 | 3.78e-09 | 0.823 | 3.84e-09 | 0.759 |
| Degree | Spatial | 0.562 | 0.428 | 1 | 0.0135 | 0.973 | -0.151 |
| Clustering | Spatial | 0.693 | -0.232 | 0.000357 | -1.02 | 8.48e-05 | -0.921 |

***Supplementary Table 6.***

**Statistical comparisons of rodent 100k neuronal culture topological fingerprint dissimilarity comparisons across generative rules.**

For each test, we quote the ANOVA p-value across the generative rules and the corresponding Cohen's *d* if the ANOVA was significant at $p<0.05$. A positive Cohen's *d* reflects that Rule A has a smaller energy than Rule B, reflecting a better fit. Generative rules have been binned across the generative model class.

| Rule A | Rule B | Glutamatergic neurons | | Motor neurons | | Dopaminergic neurons | | Cerebral organoids | |
|---|---|---|---|---|---|---|---|---|---|
| | | ANOVA p=2.03e-08 | Cohen's *d* | ANOVA p=4.79e-14 | Cohen's *d* | ANOVA p=2.93e-07 | Cohen's *d* | ANOVA p=0.000265 | Cohen's *d* |
| Homophily | Degree | 0.00652 | 0.772 | 0.0037 | 0.812 | 0.866 | 0.178 | 0.971 | -0.141 |
| Homophily | Clustering | 0.000181 | 1.8 | 0.00119 | 1.88 | 0.054 | 1.34 | 0.846 | -0.303 |
| Homophily | Spatial | 1.22e-08 | 2.07 | 3.77e-09 | 10 | 9.16e-07 | 4.34 | 0.00399 | 1.82 |
| Degree | Clustering | 0.525 | 0.248 | 0.969 | 0.0735 | 0.0838 | 0.377 | 0.959 | -0.099 |
| Degree | Spatial | 2.05e-05 | 2.25 | 3.78e-09 | 8.34 | 6.27e-07 | 4.29 | 0.000321 | 2.5 |
| Clustering | Spatial | 0.000437 | 1.83 | 3.82e-09 | 3.96 | 0.000143 | 5.48 | 0.000113 | 1.57 |

***Supplementary Table 7.***

**Statistical comparisons of human iPSC neuronal culture (DIV28 glutamatergic neurons, DIV28 motor neurons and DIV28 dopaminergic neurons) and human cerebral organoids energy comparisons across generative rules.**

For each test, we quote the ANOVA p-value across the generative rules and the corresponding Cohen's *d* if the ANOVA was significant at $p<0.05$. A positive Cohen's *d* reflects that Rule A has a smaller energy than Rule B, reflecting a better fit. Generative rules have been binned across the generative model class.

| Antibodies (human organoids) | Type | Dilution | Catalog number |
|---|---|---|---|
| Tau | Primary | 1:500 | #MN1000, ThermoFisher |
| NeuN | Primary | 1:300 | #M11954-3, Boster Bio, Pleasanton, CA, USA |
| GFAP | Primary | 1:500 | #NB300-141, Novus Biologicals, Englewood, CO, USA |
| goat anti-mouse IgG, Alexa Fluor Plus 488 | Secondary | 1:400 | #A32723, ThermoFisher |
| goat anti-rabbit IgG, Alexa Fluor 568 | Secondary | 1:400 | #A11036, ThermoFisher |
| goat anti-chicken IgY, Alexa Fluor Plus 647 | Secondary | 1:400 | #A32933, ThermoFisher |
| **Antibodies (human iPSC)** | **Type** | **Dilution** | **Catalog number** |
| mouse anti-TH | Primary | 1:500 | #MAB318, Sigma-Aldrich |
| chicken anti-MAP2 | Primary | 1:1000 | #CH22103, Neuromics (Edina, MN, USA) |
| rabbit anti-GFAP | Primary | 1:500 | #Z0334, Agilent (Santa Clara, CA, USA) |
| donkey anti-mouse 488 | Secondary | 1:250 | #A-21202, ThermoFisher |
| goat anti-chicken 647 | Secondary | 1:500 | #A-32933, ThermoFisher |
| donkey anti-rabbit 568 | Secondary | 1:250 | #A-A10042, ThermoFisher |
| **Antibodies (rodent PC)** | **Type** | **Dilution** | **Catalog number** |
| mouse anti-Synaptophysin | Primary | 1:100 | #ab8049, Abcam |
| rabbit anti-NeuN | Primary | 1:300 | #ab177487, Abcam |
| chicken anti-beta III Tubulin | Primary | 1:1000 | #ab41489 Abcam |
| donkey anti-mouse IgG, Alexa Fluor 488 | Secondary | 1:500 | #ab150105, Abcam |
| donkey anti-Chicken IgY (IgG) | Secondary | 1:400 | #703–165-155, Jackson ImmunoResearch, West Grove, USA |
| donkey anti-rabbit IgG, Alexa Fluor 405 | Secondary | 1:1000 | #ab175651, Abcam |

***Supplementary Table 8. Primary and secondary antibodies.***
